# Systematic identification of Y-chromosome gene functions in mouse spermatogenesis

**DOI:** 10.1101/2024.07.18.604063

**Authors:** Jeremie Subrini, Wazeer Varsally, Irina Balaguer Balsells, Maike Bensberg, Georgios Sioutas, Obah Ojarikre, Valdone Maciulyte, Björn Gylemo, Katharine Crawley, Katherine Courtis, Dirk G. de Rooij, James M. A. Turner

## Abstract

The mammalian Y chromosome is essential for male fertility, but how individual Y genes regulate spermatogenesis is poorly understood. To resolve this question, we generate a deletion series of the mouse Y chromosome, creating thirteen Y-deletant mouse models and conducting exhaustive reproductive phenotyping. Eight Y genes, including several that are deeply conserved and exhibit testis-specific expression, are dispensable for spermatogenesis. For others, we uncover novel functions, including a role for *Uty* in establishment and differentiation of spermatogonia, and for *Zfy2* in ensuring meiotic pairing and reciprocal recombination between the sex chromosomes. We also generate the first mouse equivalent of the human infertility AZFa deletion, revealing cumulative detrimental effects of Y-gene loss on spermatogenesis. We use single nuclei RNAseq to identify candidate mechanisms by which Y genes regulate the germ cell transcriptome and reveal an unexpected impact of Y genes on testis supporting cells. Our study represents a paradigm for the complete functional dissection of a mammalian Y chromosome and advances our knowledge of human infertility and Y-chromosome evolution.

## Main

The mammalian sex chromosomes evolved from a pair of autosomes, with the Y chromosome degenerating and ultimately losing around 92% of its ancestral gene content^1,2^. The remaining non-recombining region of the mouse Y chromosome comprises sixteen gene families, which encode proteins with predicted regulatory roles including transcription activation, ubiquitylation, chromatin modification, RNA stability and translation^1–4^ (Fig.1a). Four (*Rbmy*, *Sly*, *Srsy* and *Ssty1/2*) are ampliconic, occupying the Y-long arm and centromeric end of the Y-short arm, and are implicated in spermiogenesis and sex ratio control^5–10^. Of the remaining twelve, *Uba1y*, *Kdm5d*, *Eif2s3y*, *Uty*, *Ddx3y*, *Usp9y*, *Sry*, *Zfy1*, *Zfy2* and *Prssly* were present on the ancestral XY pair, whereas *Teyorf1* and the duplicated *H2al2y* (*H2al2b* and *H2al2c*) were more recently acquired in the mouse lineage. The majority (twelve out of sixteen) of mouse Y genes exhibit testis-biased expression. This, together with the fact that large genomic deletions on the Y chromosome are associated with fertility defects in mice and men, highlight the importance of the Y chromosome in spermatogenesis^4^. However, our understanding of the specific Y genes necessary for spermatogenesis and their precise roles remains incomplete. The functions of only a few genes have been identified. *Sry* is necessary and sufficient for testis determination^11^. *Eif2s3y* is necessary for spermatogonial proliferation, but its mechanism of action remains unknown^12,13^. *Zfy* genes have been shown to be important in meiosis and sperm morphogenesis, but their exact functions are not fully understood^14–20^. Other Y genes have been targeted with no overt fertility phenotypes, but confirmation that the targeted allele was a null was not always established, or deeper reproductive phenotyping was not performed^21–26^. For some genes, knockouts (KO) have never been made. In this study, we established a pipeline to systematically generate null Y-gene deletions and performed extensive fertility phenotyping, thereby determining which genes are involved in spermatogenesis and how they function.

**Fig. 1:**
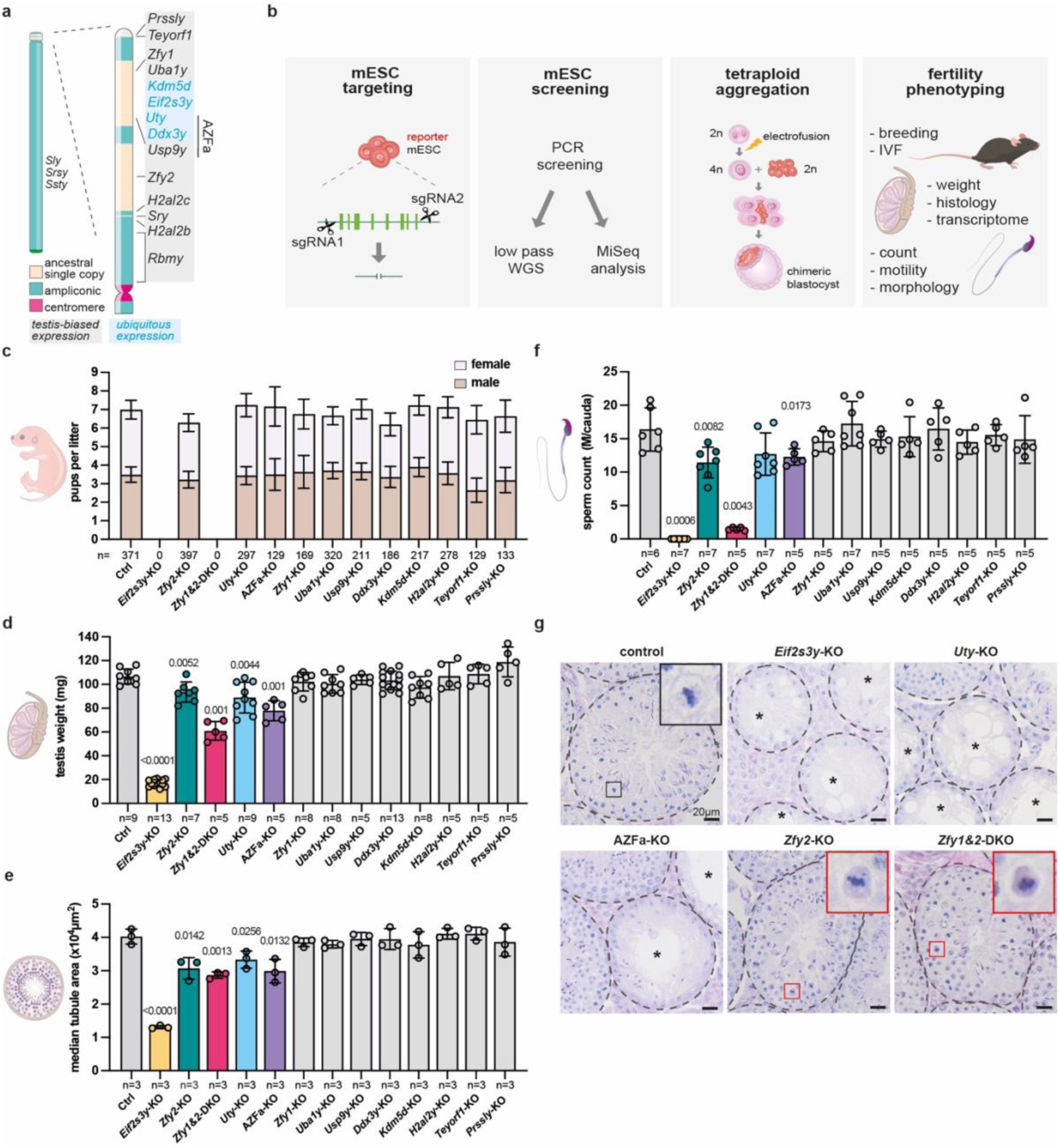
Generation and screening of thirteen Y-deletant mouse models for abnormal gamete production. **a**, Structure and gene content of the mouse Y chromosome. **b**, Experimental strategy to generate and study Y-gene deletant mouse models. **c**, Mean number of male and female pups born per litter from control and Y-deletant matings. n = total number of pups born. Error bars = 95% confidence interval. **d**, Mean testis weight in controls and Y-deletants. n = number of males. Error bars = standard deviation. **e**, Median seminiferous tubule area in control and Y-deletants. n = number of replicate males. At least 40 tubules were counted per replicate. Error bars = standard deviation. **f**, Average sperm count in controls and Y-deletants. Sperm was collected from the cauda epididymis. n = number of males. Error bars = standard deviation. **g,** PAS-stained testis sections of control, *Eif2s3y*-KO, *Uty*-KO, AZFa-KO *Zfy2*-KO and *Zfy1&2*-DKO. Tubules are circled by dotted lines. Asterisks indicate tubules with severe germ cell depletion. Black inset: control metaphase, red insets: dark cytoplasmic signal is visible, indicative of apoptosis. Statistical analysis by two-sided Chi-Square test for **c**, Mann Whitney test for **d** and **f** and two tailed t-test for e. Significant p values (<0.05) are shown.

### Generating mouse models with Y-gene deletions

We targeted Y genes in XY mouse embryonic stem cells (mESCs), which were then used to produce Y-KO animals. For each Y gene, the entire coding region was excised using CRISPR-Cas9 with two flanking sgRNAs, thereby generating null alleles (Fig.1b, Extended data Fig.1a, b). The exception was the duplicated *H2al2y*, for which we used one sgRNA to generate indels in both copies. We used this approach for *H2al2y* because the two identical copies *H2al2b* and *H2al2c* flank *Sry*, meaning that a whole-gene deletion strategy could inadvertently remove *Sry* and cause sex reversal (Fig.1a, Extended data Fig.1c). For *Zfy1* and *Zfy2*, we created individual deletions and a combined *Zfy1&2*-double knockout (DKO) deletion, allowing us to examine functional overlap between these paralogues. We also generated the first animal model of the *AZ*oospermia *F*actor a (AZFa) deletion, encompassing *Usp9y*, *Uty* and *Ddx3y,* which in men causes the most severe form of infertility characterised by a complete absence of germ cells^27^. We used this model to understand the aetiology of AZFa-related infertility, and to examine functional divergence of the Y chromosome in mice and men. The multi-gene KOs also allowed us to interrogate combinatorial effects of Y-gene loss on spermatogenesis.

Once fully validated by low-pass whole genome sequencing (Extended data Fig.1e, f) and MiSeq analysis of on- and off-target mutations (Extended data Fig.1g, h), deletant mESC lines were used to make animals. This is usually achieved by blastocyst mESC injection, and breeding of resulting chimeras to attain germline transmission. Such an approach was not suitable for mutations that could compromise fertility. We therefore used tetraploid aggregation to generate founders that were fully derived from the mESCs, i.e., were fully mutant. We improved the typically low birth and offspring survival rates associated with this technology ^28–31^ (see Methods for details) and generated founders from all our Y-KO mESC lines. ^28–31^ We conclude that *Eif2s3y, Zfy1, Zfy2*, *Uty*, *Uba1y*, *Usp9y, Ddx3y*, *Kdm5d*, *H2al2y, Teyorf1*, and *Prssly* are all dispensable for embryonic survival.

### Spermatogenic defects in a subset of Y-gene KOs

To determine which Y genes are necessary for reproduction, Y-KO males were mated with wild type females. Consistent with previous studies, *Eif2s3y*-KO and *Zfy1&2*-DKO founders were infertile^15,32^ (Fig.1c). The remaining eleven Y-deletant lines all produced offspring (Fig.1c). Thus, three of the eleven deleted Y gene families are necessary but not sufficient for mice to father offspring^11–13,17^: *Sry* (necessary for testis determination), *Eif2s3y* (necessary for spermatogonial proliferation), and *Zfy* (at least one copy needed to produce functional sperm).

Ability to reproduce does not necessarily mean that spermatogenesis is normal. We therefore developed a phenotyping pipeline to exhaustively characterise the reproductive fitness of individual Y-deletants. We assayed litter sizes, offspring sex ratios, testis weights and histology, sperm counts, sperm head morphology, sperm motility, and *in vitro* fertilization (IVF) success rates. Notably, all these parameters remained unaffected for deletion of *Zfy1*, *Uba1y*, *Usp9y*, *Ddx3y*, *Kdm5d*, *H2al2y*, *Teyorf1* and *Prssly,* showing that these genes are dispensable for mouse spermatogenesis (Fig.1c-f, Fig. 2a,d-f, Extended data Fig.2a-i).

**Fig. 2:**
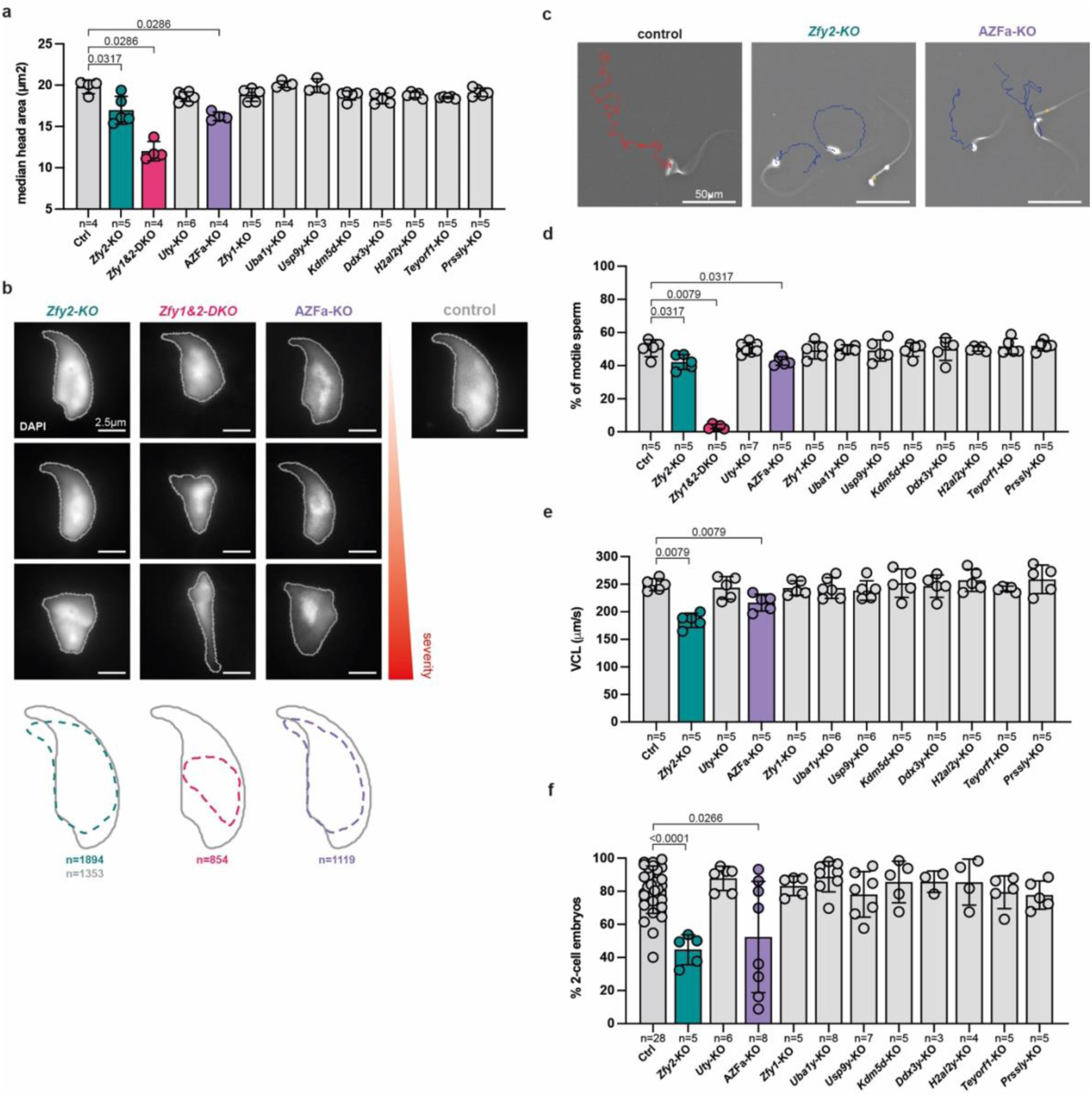
Screening for reduced gamete quality in thirteen Y-deletant mouse models. **a**, Median sperm head area for control and Y-deletants. n = number of males. Error bars = standard deviation. **b**, DAPI-stained sperm heads from *Zfy2*-KO, *Zfy1&2*-DKO and AZFa-KO showing a range of abnormalities, from least (top) to most severe (bottom), compared to a control sperm head with stereotypical shape. The average sperm profile of the three deletants is compared to that of the control. n = number of sperm heads used to build the average outline. **c**, Sperm from control, *Zfy2*-KO and AZFa-KO tracked over one second. Red trajectory represents a rapid progressive sperm and blue trajectories slow, non-progressive sperm. **d**, Average percentage of motile sperm in the cauda epididymis of controls and Y-deletants. n = number of males. Error bars = standard error of the mean. **e**, Average sperm curvilinear velocity in controls and Y-deletant. n = number of males. Error bars = standard error of the mean. **f**, Average fertilisation rate of sperm from control and Y-deletant males in IVF assays. n = number of males. Error bars = standard deviation. Statistical analysis by Mann Whitney test, with significant p values (<0.05) shown.

In contrast, *Eif2s3y*-KO, *Uty*-KO, AZFa-KO, *Zfy2*-KO, and *Zfy1&2*-DKO males all exhibited defective germ cell production, manifested as reduced testis weight, seminiferous tubule area, and sperm counts (Fig.1d-f). We found Y genes to be involved in all the major steps of spermatogenesis: mitosis, meiosis and spermiogenesis. *Eif2s3y-*KO males showed severe germ cell depletion, consistent with the role of this gene in spermatogonial proliferation^12^ (Fig.1g). Interestingly, *Uty*-KO and AZFa-KO males also exhibited germ cell-depleted tubules, revealing a role for *Uty* in spermatogonia (Fig.1g, Fig.3c, Extended data Fig.3a; and see later).

**Fig. 3:**
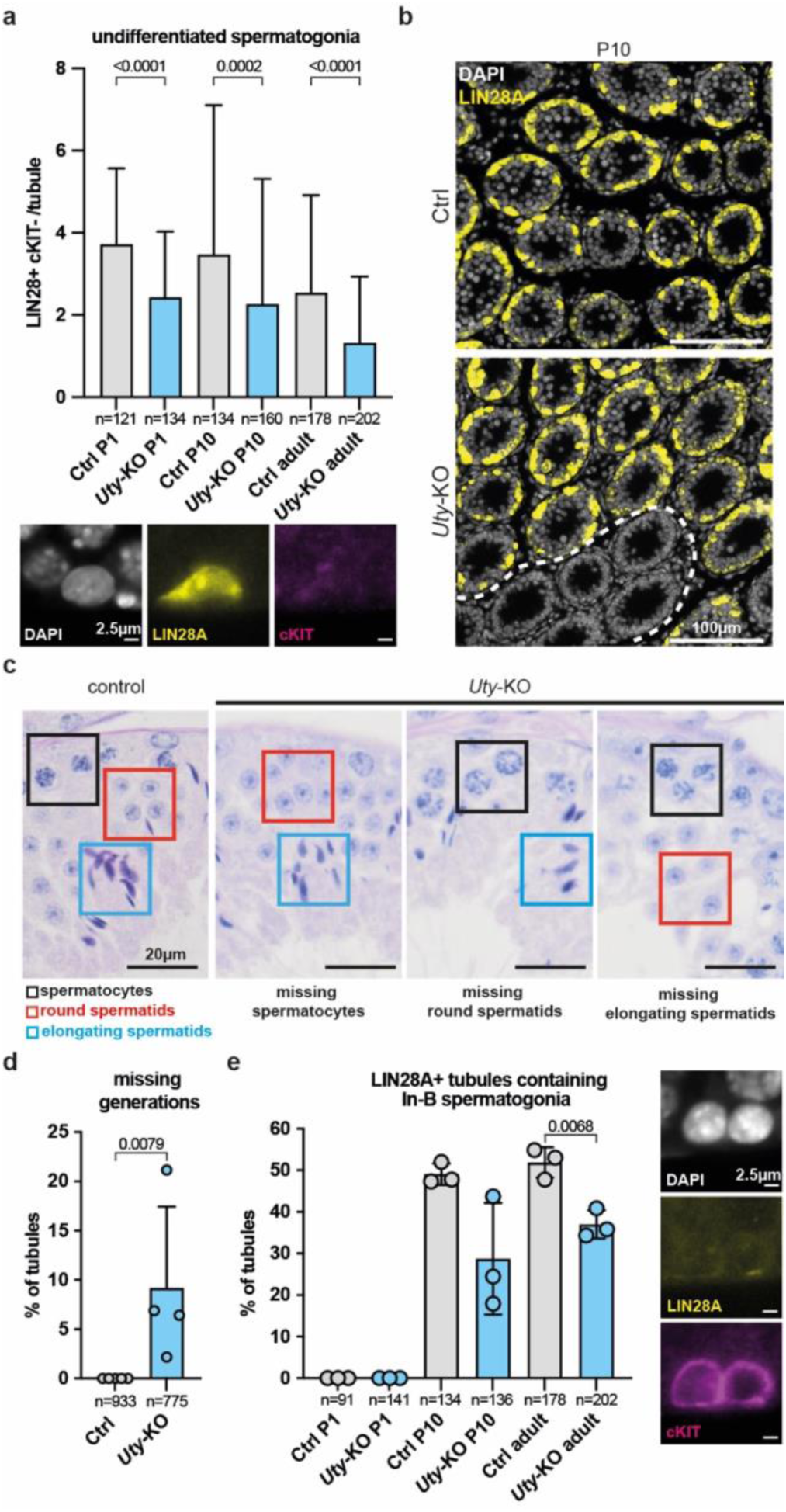
Defects in the establishment and differentiation of the spermatogonial pool in *Uty*-deleted mice. **a**, Number of undifferentiated spermatogonia per tubule in control and *Uty*-KO P1, P10 and adults, quantified in testis immunostaining for LIN28A and cKIT. n = number of tubules counted across three biological replicates. p values calculated by Mann Whitney test. **b**, P10 testis sections immunostained for LIN28A. Dashed lines encircle tubules with no germ cells. **c**, PAS-stained testis sections, with *Uty*-KO exhibiting different missing generations compared to control. **d**, Percentage of tubules exhibiting missing generations. n = number of tubules counted across four biological replicates. **e**, Quantification of percentage of tubules with differentiating In-B spermatogonia using P1, P10 and adult testis sections immunostained with LIN28A and cKIT. Tubules lacking progenitor cells (with no LIN28A+ cells) were excluded. n = number of tubules counted across three biological replicates. All error bars = standard deviation. Statistical analysis by Mann Whitney test for **a** and **d** and two tailed t-test for **e**. Significant p values (<0.05) are shown.

*Zfy2*-KO and *Zfy1&2*-DKO males exhibited phenotypes at two stages of spermatogenesis. The first, not previously described, occurred during meiosis, with some metaphase cells undergoing apoptosis (Fig.1g; and see later). The second occurred during spermiogenesis, with abnormal sperm morphology and motility affecting fertilisation (Fig.2a-f, Extended data Fig.2b-i, Extended data Fig.3b-d). These defects were more severe in the *Zfy1&2*-DKO males, illustrating partially redundant roles of *Zfy1* and *Zfy2* in developing sperm (Fig.1c, Fig.2, Extended data Fig.3b-d).

Notably, AZFa KO males also exhibited abnormal sperm morphology and motility, ultimately causing decreased IVF success (Fig.2a-f). Sperm defects were not observed in *Uty*, *Usp9y* or *Ddx3y* single mutants (Extended data Fig.2a-i), indicating that multi-Y gene deletions can create novel phenotypes that are not observed in single gene deletants and unmask potential synergistic roles of Y genes. We conclude that genes exhibiting no observable effects when deleted individually may still impact spermatogenesis. Furthermore, the observation that, unlike men, AZFa-KO mice can produce sperm, provides the first experimental evidence that the mouse and human Y chromosomes are divergent in their spermatogenic functions.

### *Uty* regulates spermatogonial pool establishment and differentiation

Adult *Uty*-KO males exhibited some tubules with few or no germ cells, suggesting a defect at the spermatogonial stage (Fig.1g). This phenotype could arise through failure to establish, differentiate or renew the spermatogonial stem cell (SSC) pool. There was no significant difference in testis weights, seminiferous tubule area, or sperm counts between adult (13-15 weeks) and aged (43 weeks) *Uty*-KO males, suggesting that SSC self-renewal was unaffected (Extended data Fig.4a-c). We therefore focused on SSC establishment and spermatogonial differentiation. To examine SSC establishment, testis sections were co-stained for LIN28A (a marker of undifferentiated and A1-A4 spermatogonia), and cKIT (a marker of differentiating A1 to B spermatogonia) at postnatal day (P)1, P10 and adult ^33,34^. The number of undifferentiated spermatogonia (LIN28A+; cKIT-) was significantly reduced in *Uty*-KO males at all ages (Fig.3a-b). Accordingly, the number of differentiating A1-A4 and In-B spermatogonia (cKIT+; LIN28A-) was also lower (Extended data Fig.4d-e). The finding that spermatogonia were reduced as early as P1 demonstrates that *Uty* is required for the proper establishment of the spermatogonial pool.

To examine whether differentiation was also compromised, we examined testis histology and the proportion of undifferentiated versus differentiated spermatogonia in *Uty*-KO males. Remarkably, some tubules were missing whole generations of germ cells. Because this defect was not cell-type specific and affected spermatocytes, round, and elongating spermatids at different stages of the seminiferous cycle, it was most likely caused by an early differentiation failure, in spermatogonia (Fig.3c, d). Supporting this hypothesis, we observed a reduction in the proportion of tubules containing late-stage differentiating spermatogonia (cKIT+; LIN28A-) and in the ratios of In-B to A1-A4 spermatogonia (Fig.3e, Extended data Fig.4f). We conclude that *Uty* has two functions in early spermatogenesis, first regulating both spermatogonial stem cell establishment, and later spermatogonial differentiation.

### *Zfy2* regulates X-Y chromosome pairing at meiosis

In *Zfy2*-KO and *Zfy1&2*-DKO males, we observed apoptosis in meiotic metaphases (Fig.1g), which we quantified by cPARP immunostaining (Fig.4a). Metaphase apoptosis is commonly caused by misaligned chromosomes, which trigger the spindle assembly checkpoint (SAC)^35^. Indeed, misaligned chromosomes were clearly apparent in DAPI-stained *Zfy2*-KO and *Zfy1&2*-DKO metaphase cells (Fig.4a-c). Since sex chromosomes are more prone to misalignment than autosomes^35^, we used X and Y chromosome painting to determine whether they are misaligned. In 99.51% of cases, the misaligned chromosome was the X or the Y, confirming a sex chromosome-specific effect (Fig.4c). Despite this X-Y misalignment, there was no increase in sex chromosome aneuploid spermatids in *Zfy2*-deficient males (Extended data Fig.5a). This finding indicated that, contrary to a previous report^18^, the SAC functions efficiently without *Zfy2* (see Supplementary Discussion).

**Fig. 4:**
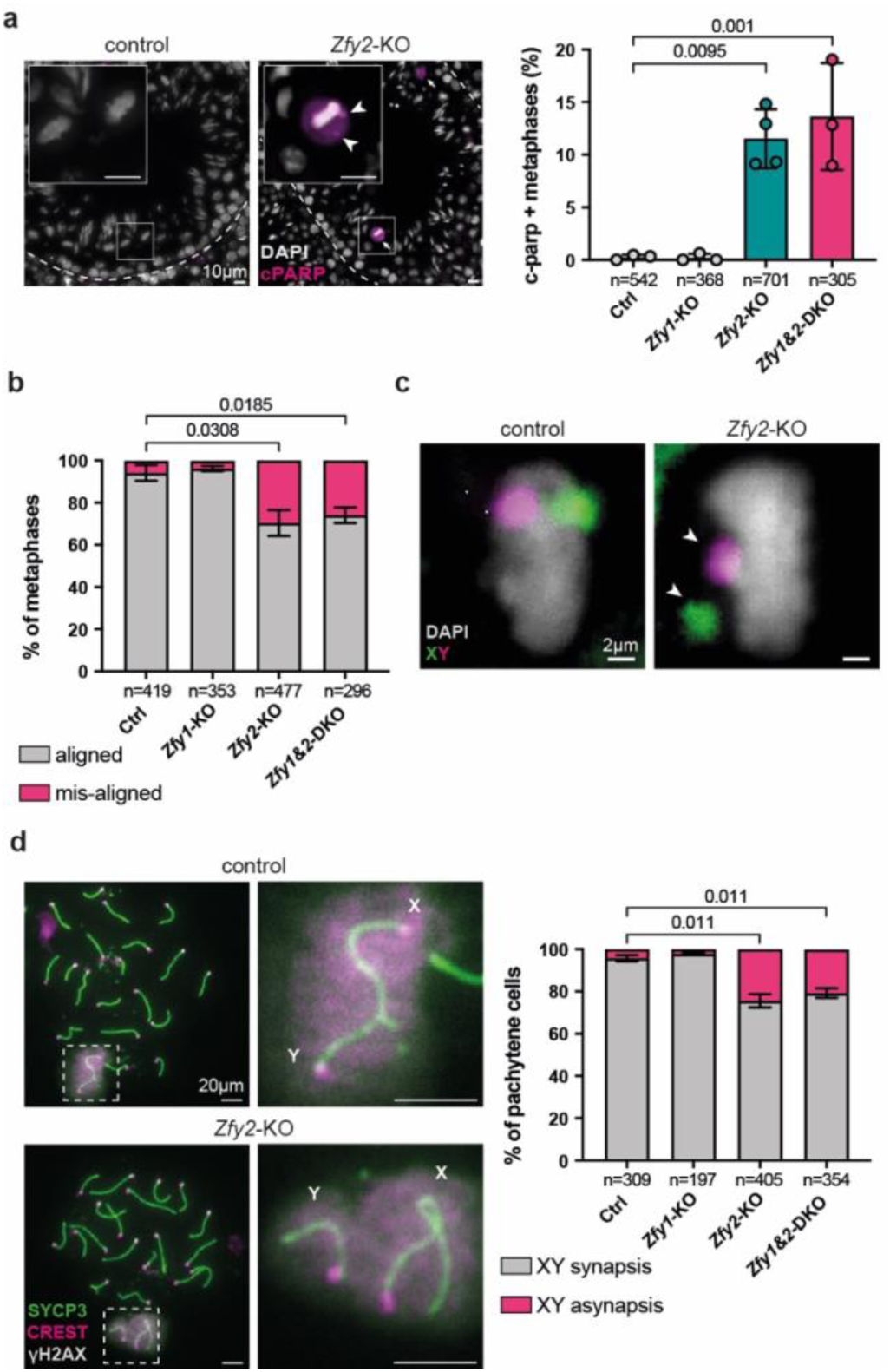
Defects in meiotic sex chromosome pairing in the absence of Zfy2. **a**, cPARP immunostaining in testis sections to quantify metaphase apoptosis in control and *Zfy* deletants. Arrows indicate apoptotic metaphase cells. Arrowheads point to misaligned chromosomes. Insets show example metaphase cells. n = number of metaphases counted across three-four replicates. Error bars = standard deviation. **b**, Percentage of metaphase cells with aligned and misaligned chromosomes in control and *Zfy* deletants. n = number of metaphases counted across three biological replicates. Error bars = standard error of the mean. **c**, Representative aligned and misaligned metaphase plates in testis sections painted with X (green) and Y (magenta) whole chromosome paints. Arrowheads point to misaligned chromosomes. **d**, Pachytene spermatocytes immunostained for SYCP3 (green), CREST (magenta) and γH2AX (grey) to quantify X-Y asynapsis in control and *Zfy* deletants. Insets show sex chromosomes, with synapsed and asynapsed X-Y in control and *Zfy2*-KO, respectively. n = number of pachytene cells counted across three biological replicates. Error bars = standard error of the mean. Statistical analysis by unpaired two-tailed t test, with significant p values (<0.05) shown.

We suspected that the misalignment phenotype at metaphase resulted from a failure to establish synapsis and recombination between the X and Y chromosomes earlier, at pachynema. To test this hypothesis, we quantified synapsis using antibodies to the axial elements (SYCP3), centromeres (CREST), and the X-Y pair (γH2AX) (Fig.4d). *Zfy2*-KO and *Zfy1&2*-DKO males exhibited a higher frequency of X-Y asynapsis than controls, while autosomal synapsis was unaffected (Fig.4d; Extended data Fig.5b). The incidence of X-Y asynapsis was similar between *Zfy2*-KO and *Zfy1&2*-DKO males, and *Zfy1*-KOs were unaffected, indicating a specific role for *Zfy2* and not *Zfy1* in promoting X-Y pairing. Asynapsed X-Y pairs failed to form crossovers, as demonstrated by MLH3 staining (Extended data Fig.5c). ZFY2 thus promotes X-Y synapsis and reciprocal recombination.

### The impact of Y genes on the testis transcriptome

To interrogate how Y genes impact the testis transcriptome, we performed bulk RNAseq on adult testes from our thirteen mutants. Consistent with their more dramatic spermatogenic defects, *Eif2s3y*-KO, *Zfy1&2*-DKO, and *Zfy2*-KOs showed the greatest separation from other samples on a Principal Component Analysis (PCA; Extended data Fig.6a, b), and exhibited the highest number of differentially expressed (DE) genes compared to controls (Fig.5a). Gene set enrichment analysis (GSEA) revealed that gene ontology terms related to spermatogenesis were deregulated in these mutants (Fig.5a, Extended data Fig.6c, d; Supplementary Table 1 and 2). In mutants with no phenotypes, we observed a surprisingly wide range of effects on gene expression. While some mutants exhibited few or no DE genes (e.g., *Teyorf1*-KO), others exhibited hundreds of DE genes (e.g., *Ddx3y*-KO and *Kdm5d*-KO) (Fig.5a, Supplementary Table 1 and 2). In the case of *Ddx3y*-KO other genes associated with RNA helicase activity were also downregulated (Extended data Fig.6e). We conclude that Y genes for which deletion creates no apparent spermatogenic phenotype can still regulate multiple downstream targets in the testis.

**Fig. 5:**
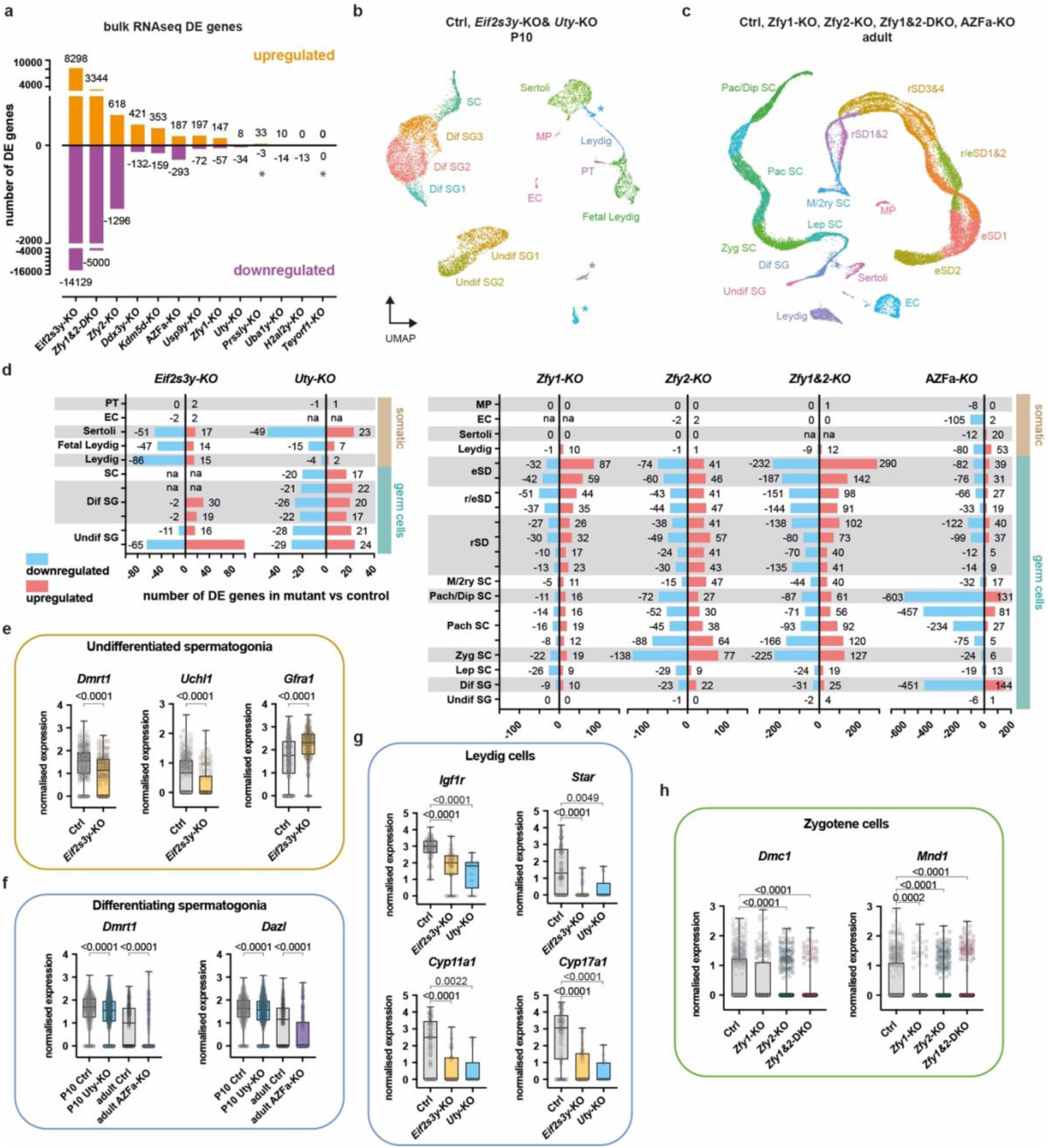
Transcriptional deregulation in Y-deletant testes. **a**, Quantification of differentially expressed (DE) genes in bulk RNAseq of Y-KO testes compared to controls. A threshold of Log2 fold change ≥0.5 was used. Asterisks indicate Y-genes not annotated in the mouse genome assembly which thus won’t appear as downregulated. **b-c**, Uniform manifold approximation and projection (UMAP) of P10 and adult single nuclei RNAseq (snRNAseq) datasets showing undifferentiated and differentiating (Undif and Dif) spermatogonia (SG); leptotene (Lep); zygotene (Zyg); pachytene (Pach); diplotene (Dip); metaphase (M) and secondary (2ry) spermatocytes (SC); round and elongating spermatids (rSD and eSD); foetal Leydig; adult Leydig; Sertoli; peritubular (PT); endothelial (EC) and macrophage (MP) cells. Asterisks show unknown clusters with no clear signatures. **d**, Quantification of DE genes for each cell type in P10 *Eif2s3y* and *Uty* KOs and adult *Zfy1*, *Zfy2*, *Zfy1&2* and AZFa KOs. Cell types with less than 10 nuclei in the mutants are marked as na. **e**, Expression of the differentiation markers *Dmrt1*, *Uchl1* and the progenitor marker *Gfra1* in *Eif2s3y*-KO and control Undif SG1. **f**, Expression of the differentiation markers *Dmrt1* and *Dazl* in *Uty*-KO, AZFa-KO and control Dif SG. **g**, Expression of members of the steroid synthesis pathway in *Eif2s3y*-KO, *Uty*-KO and control Leydig cells. **h**, Expression of the recombinases *Dmc1* and *Mnd1* in *Zfy1*, *Zfy2*, *Zfy1&2* KOs and control zygotene cells. Statistical analysis by Kolmogorov Smirnov test, with significant p values (<0.05) shown.

To gain deeper molecular insights into each spermatogenic defect, we performed single nuclei RNA sequencing on *Eif2s3y*-KO, *Uty*-KO, AZFa-KO, *Zfy2*-KO, and *Zfy1&2*-DKO males. P10 testes from *Eif2s3y* and *Uty*-KOs were used to enrich for spermatogonia, since this was where the phenotypes manifested, and adult testes were used for *Zfy1*, *Zfy2*, *Zfy1&2*, and AZFa-KOs to capture all cell types (Fig.5b, c). Using published marker genes ^36,37^, we identified all the expected somatic testis and germ cell populations (Fig.5b-c, Extended data Fig.7a-c, Extended data Fig.8a-c).

By quantifying DE genes across clusters, we identified transcriptomic deregulation in unexpected cell types (Fig.5d, Supplementary Tables 3 to 8). Interestingly, in *Eif2s3y*-KO, the earliest undifferentiated spermatogonial cluster had the most DE genes (Fig.5d). The transcriptional deregulation therefore begins much earlier than the known proliferation defect, which manifests in differentiating spermatogonia^12,32,38^. Moreover, *Eif2s3y*-KOs, *Uty*-KOs and AZFa-KOs had extensive deregulation in Leydig and Sertoli cells, revealing a surprising impact on the somatic testis transcriptome (Fig.5d). Notably, despite *Zfy2* being predominantly expressed in spermatids^16^, the highest number of DE genes in the *Zfy2*-KO was found at zygonema (Fig.5d). Strikingly, in AZFa-KOs, the number of DE genes increased throughout pachynema, even though no visible defects were observed at this stage (Fig.5d).

Using gene set enrichment analysis, we identified multiple pathways affected by Y-gene loss that may contribute to the spermatogonial defects in *Eif2s3y*, *Uty*, and AZFa deletants, the meiotic defects in *Zfy2* and *Zfy1&2* mutants, and the spermatid defects in *Zfy2*, *Zfy1&2*, and AZFa mutants (Supplementary Tables 9 to 14). To find candidate genes contributing to each phenotype, we also examined DE genes along with the expression of known spermatogenic regulators. In *Eif2s3y*-KOs, genes involved in spermatogonial differentiation, including *Dmrt1*, *Uchl1* and *Stra8* were downregulated, while markers of the undifferentiated state, including *Gfra1* and *Etv5* were upregulated (Fig.5e, Extended data Fig.7d, e). *Eif2s3y* could therefore act to suppress the program that maintains an undifferentiated state^39^, or it could promote activation of the differentiation program. In *Uty*-KOs, known regulators of spermatogonial development were downregulated, including *Dmrt1* and *Dazl* (Fig.5f). Strikingly, the steroidogenesis pathway, including genes such as *Igf1r*, *Star*, *Cyp11a1* and *Cyp17a1*, was downregulated in Leydig cells of *Eif2s3y*, *Uty* and AZFa deletants. Since the production of steroid hormones like testosterone is crucial for germ cell homeostasis^40^, this finding further emphasise the potential contribution of somatic cell deregulation to germ cell phenotypes (Fig.5g, Extended data Fig.7f, g, Extended data Fig.8d).

Interestingly, we also found that multiple factors regulating meiotic synapsis and recombination, including *Dmc1*, *Mnd1* and *Rad51ap2* were downregulated in *Zfy2*-deficient testes, implicating these factors in the X-Y pairing phenotype (Fig.5h, Extended data Fig.8e). Comparing DE genes in all our *Zfy* deletants allowed us to identify putative stage-specific shared and unique targets of ZFY1 and ZFY2 transcription factors (Extended data Fig.8f-g, Supplementary Table 15). *Zfy1&2*-DKOs shared a higher number of DE genes with *Zfy2* than with *Zfy1* mutants, further indicating that *Zfy2* is the main contributor to the *Zfy1&2*-DKO phenotypes. Surprisingly however, most DE genes in the *Zfy1&2*-DKOs were not shared with either *Zfy1* or *Zfy2* mutants. This indicated that the effects of deleting both *Zfy* genes are not merely additive, but also combinatorial.

## Discussion

Here, we generate a deletion series for the mouse Y chromosome and use an exhaustive reproductive phenotyping pipeline to provide a comprehensive overview of Y-gene functions in spermatogenesis. Strikingly, we reveal that multiple Y genes, several of which are deeply conserved and exhibit testis-biased expression, are dispensable for the germline. However, we also demonstrate that more Y genes are required for normal murine spermatogenesis than previously known, and function in more spermatogenic processes than previously thought. We also generate the first transcriptomic atlas of Y-mutant testes, revealing how Y genes regulate testis gene expression and providing a resource to the research community.

Our study provides new perspectives into human infertility. Most known cases of human Y chromosome-related infertility are caused by either AZFa, AZFb or AZFc deletions^27^. Since these deletions remove multiple Y genes, identifying which is causative remains a challenge in clinical genetics. Even though Y deletions may not always cause the same phenotypes in mice and humans^41^, our work raises the possibility that the aetiology of some infertility cases may result from the cumulative loss of several Y genes, each of which only having a minimal impact on spermatogenesis.

While most Y-genes have been lost during evolution, cross-species comparisons have revealed that many that persist are under strong purifying and positive selection and are therefore likely to retain critical functions^1,2,42^. Our finding that most Y genes are dispensable for spermatogenesis may therefore seem surprising. If some mouse Y genes are indeed non-functional, they most likely lost their utility relatively recently. Another possibility is that these genes confer a slight yet significant reproductive fitness advantage that was not captured by our study. These minor roles could be remnants of previously more prominent functions, which might now be shared with autosomal or X-linked homologues. An alternative is that these genes have been retained because they perform functions in non-reproductive organs. A rapidly increasing volume of research supports this hypothesis, linking Y chromosome loss to ageing and a wide range of conditions including cancer and heart disease^43–48^. Our Y-deletant mouse models present a valuable resource with which to unravel these phenotypes at the single gene level.

---

**Extended data Fig. 1:**
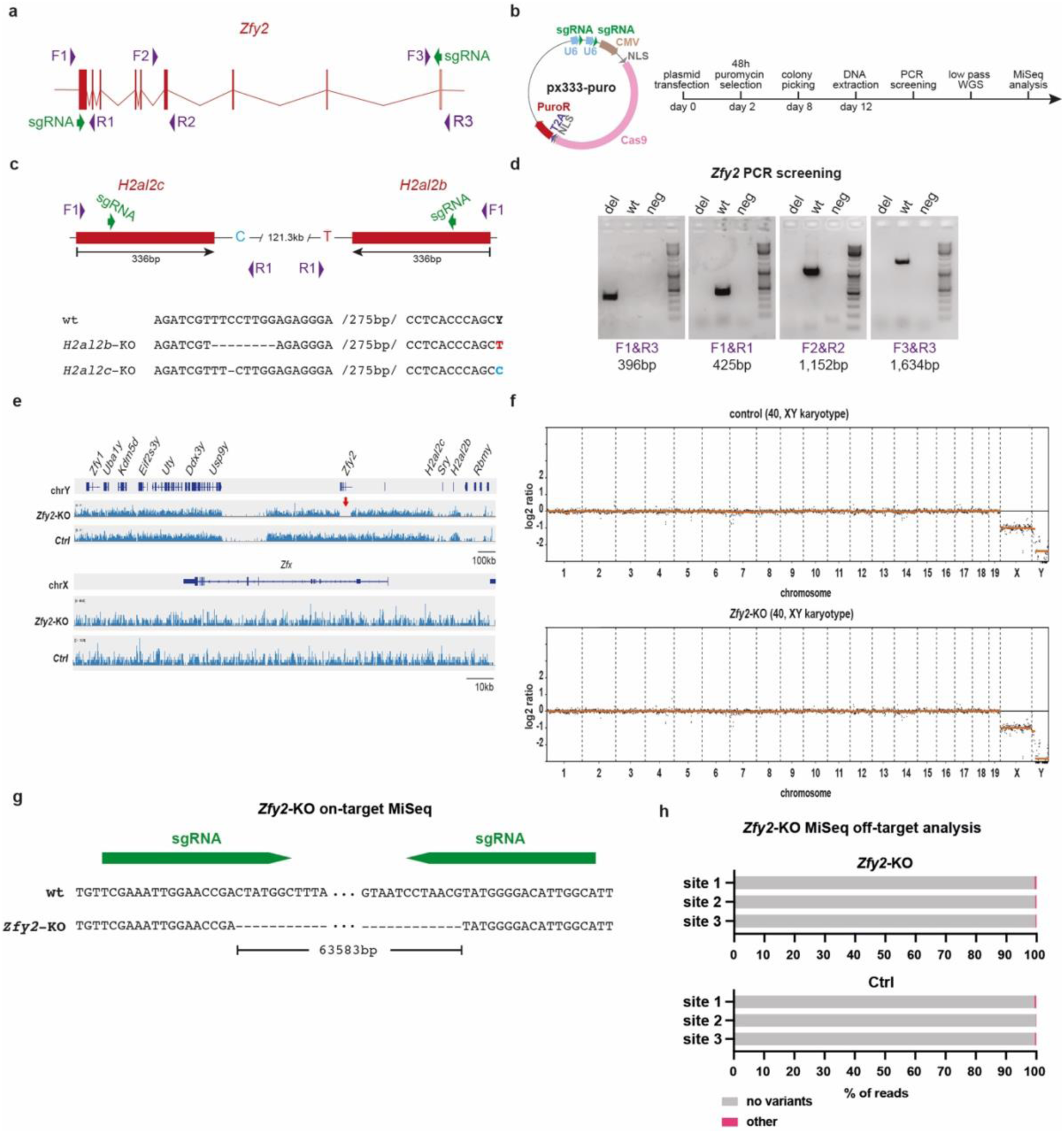
Y-gene targeting and screening pipeline in mouse embryonic stem cells. **a**, The *Zfy2* locus showing the location of sgRNAs and primers as an example of single gene targeting strategy. **b**, Map of the “all in one” px333-puro plasmid and timeline of mESC targeting. **c**, The duplicated *H2al2c* and *H2al2b* loci showing the sgRNA, primers and SNP used for targeting and sequencing. MiSeq reads of a clone with successful editing in both copies are shown. **d**, PCR screening approach for *Zfy2* targeting. Deletion, upstream boundary, exonic and downstream boundary PCRs are shown (see panel **a**, for reference primers) for a *Zfy2*-KO clone, wildtype control, and negative control (water). **e**, Low pass whole-genome sequencing (WGS) reads for *Zfy2*-KO and control clones mapped to the short arm of the Y chromosome and *Zfx*. Red arrow indicates successful deletion of *Zfy2*. **f**, Chromosomal copy number analysis for a control and *Zfy2*-KO clone showing normal karyotype. Analysis performed using the QDNAseq R package on a low pass WGS dataset. **g**, MiSeq reads of a *Zfy2*-KO clone showing a 63,583bp deletion of the *Zfy2* locus. **h**, Analysis of MiSeq reads of a *Zfy2*-KO and control wildtype clone at three potential off-target sites. Analysis performed with the CrispRVariants R package.

**Extended data Fig. 2:**
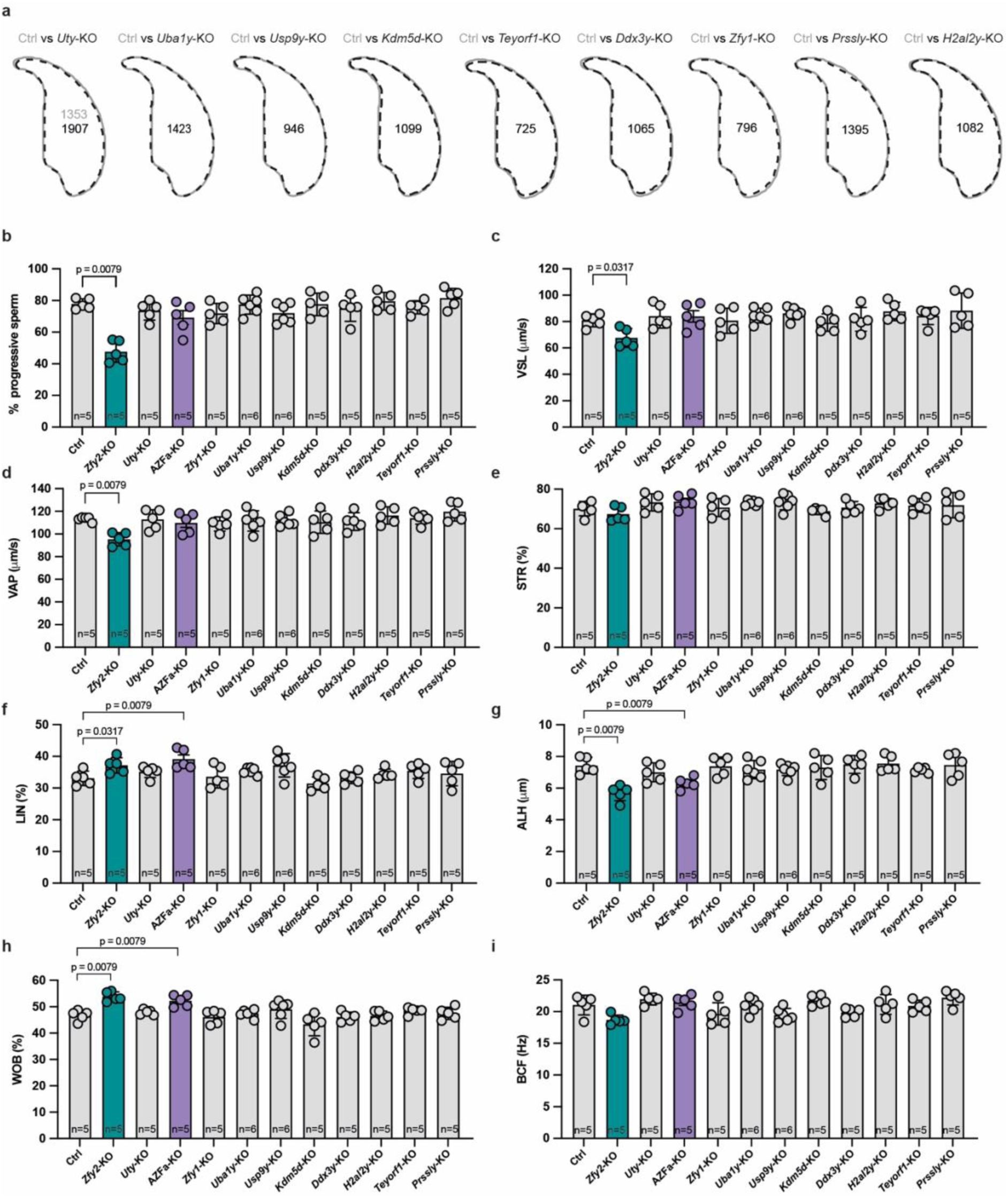
Sperm head morphology and motility analysis in Y-gene deletants. **a**, Average sperm head profile compared between control and Y deletants showing no abnormalities. The number of sperm heads used to build the consensus morphologies are indicated. **b**, Average percentage of progressive sperm for control and Y deletant males. **c**, Average straight-line velocity (VSL) of sperm from control and Y deletant males. **d**, Average path velocity (VAP) of sperm from control and Y deletant males. **e**, Mean straightness (STR) of sperm from control and Y deletant males. **f**, Mean linearity (LIN) of sperm from control and Y deletant males. **g**, Mean lateral head displacement (ALH) of sperm from control and Y deletant males. **h**, Mean wobble (WOB) of sperm from control and Y deletant males. **i**, Mean beat cross frequency (BCF) of sperm from control and Y deletant males. All n = number of males. All Error bars = standard error of the mean. All Statistical analysis by Mann Whitney test.

**Extended data Fig. 3:**
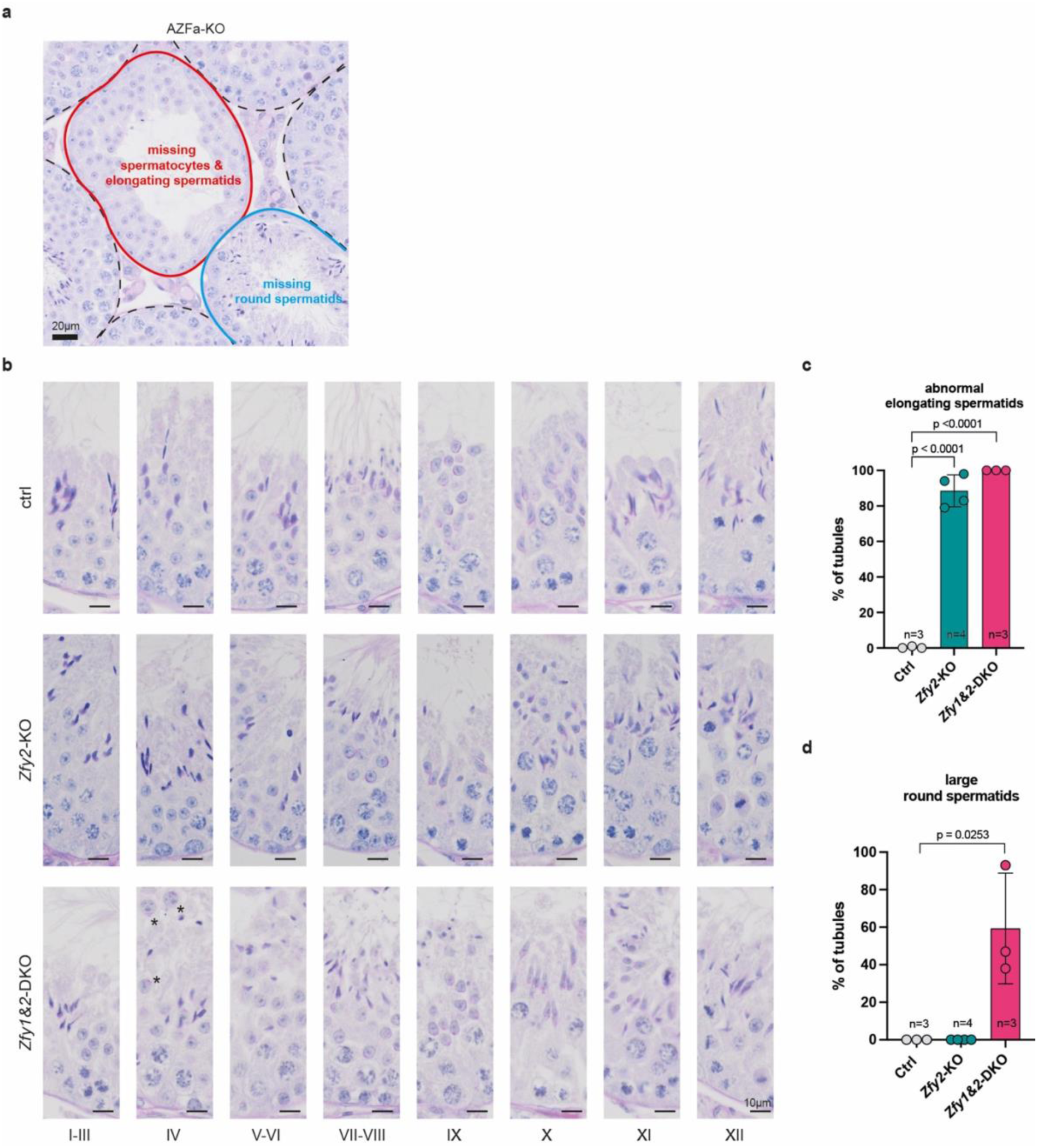
Abnormal seminiferous cycle and tubules in *Eif2s3y* and *Zfy* deletant models. **a**, PAS-stained testis section of AZFa-KO showing tubules with missing generations (red and blue outline) compared to normal tubules (black dashed outline). **b**, PAS-stained testis sections from control, *Zfy2*-KO and *Zfy1&2*-DKO at different stages of the seminiferous cycle. Asterisks mark “large”, likely diploid, round spermatids. **c**, Percentage of seminiferous tubules containing more than five elongating spermatids with abnormal morphology in controls, *Zfy2*-KO and *Zfy1&2*-DKO. **d**, Percentage of seminiferous tubules containing more than five “large”, likely diploid, round spermatids in controls, *Zfy2*-KO and *Zfy1&2*-DKO. All n = number of males. All Error bars = standard deviation. All Statistical analysis by unpaired two tailed t test.

**Extended data Fig. 4:**
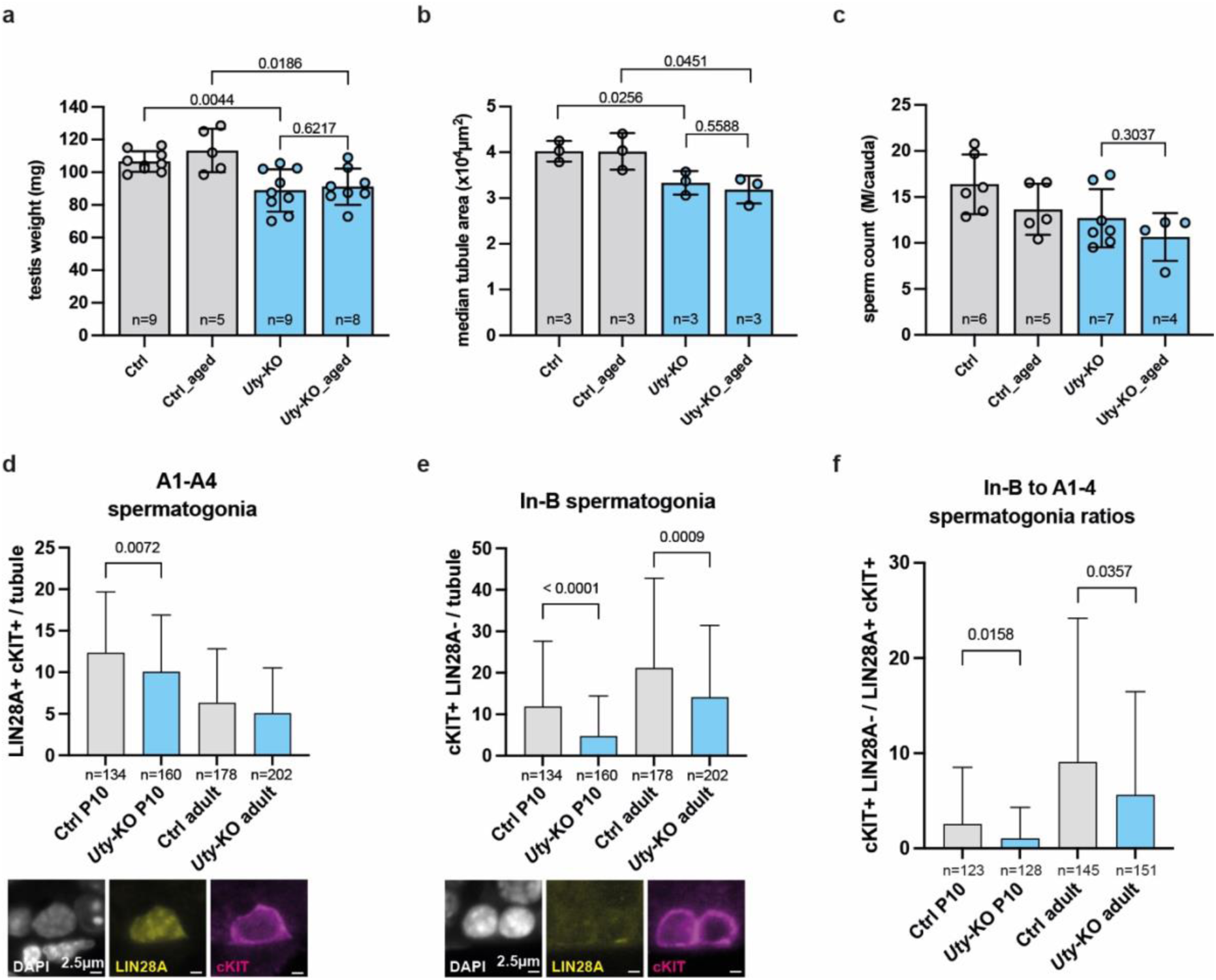
Characterisation of *Uty*-KO spermatogenic defects in juvenile, adult, and aged males. **a**, Average testis weight of adult (13-15 weeks) and aged (43 weeks) controls and *Uty*-KO. n = number of males. Statistical analysis by Mann Whitney test. **b**, Median seminiferous tubule area of adult and aged controls and *Uty*-KO control. n = number of replicate males. At least 40 tubules were counted per replicate. Statistical analysis by unpaired two tailed t test. **c**, Average cauda epididymis sperm count per animal for adult and aged controls and *Uty*-KO. n = number of males. Statistical analysis by unpaired two tailed t test. **d**, Quantification of A1-A4 spermatogonia per tubule in P10 and adult controls compared to *Uty*-KO mice, identified by positive immunostaining for LIN28A and cKIT. n = number of tubules across three biological replicates. Statistical analysis by Mann Whitney test. **e**, Quantification of In-B spermatogonia per tubule in P10 and adult controls compared to *Uty*-KO mice, identified by positive cKIT signal and negative LIN28A staining. n = number of tubules across three biological replicates. Statistical analysis by Mann Whitney test. **f**, Quantification of In-B to A1-A4 spermatogonia ratios between controls and *Uty*-KO in P10 and adult tubules. n = number of tubules. Statistical analysis by Mann Whitney test. All Error bars = standard deviation.

**Extended data Fig. 5:**
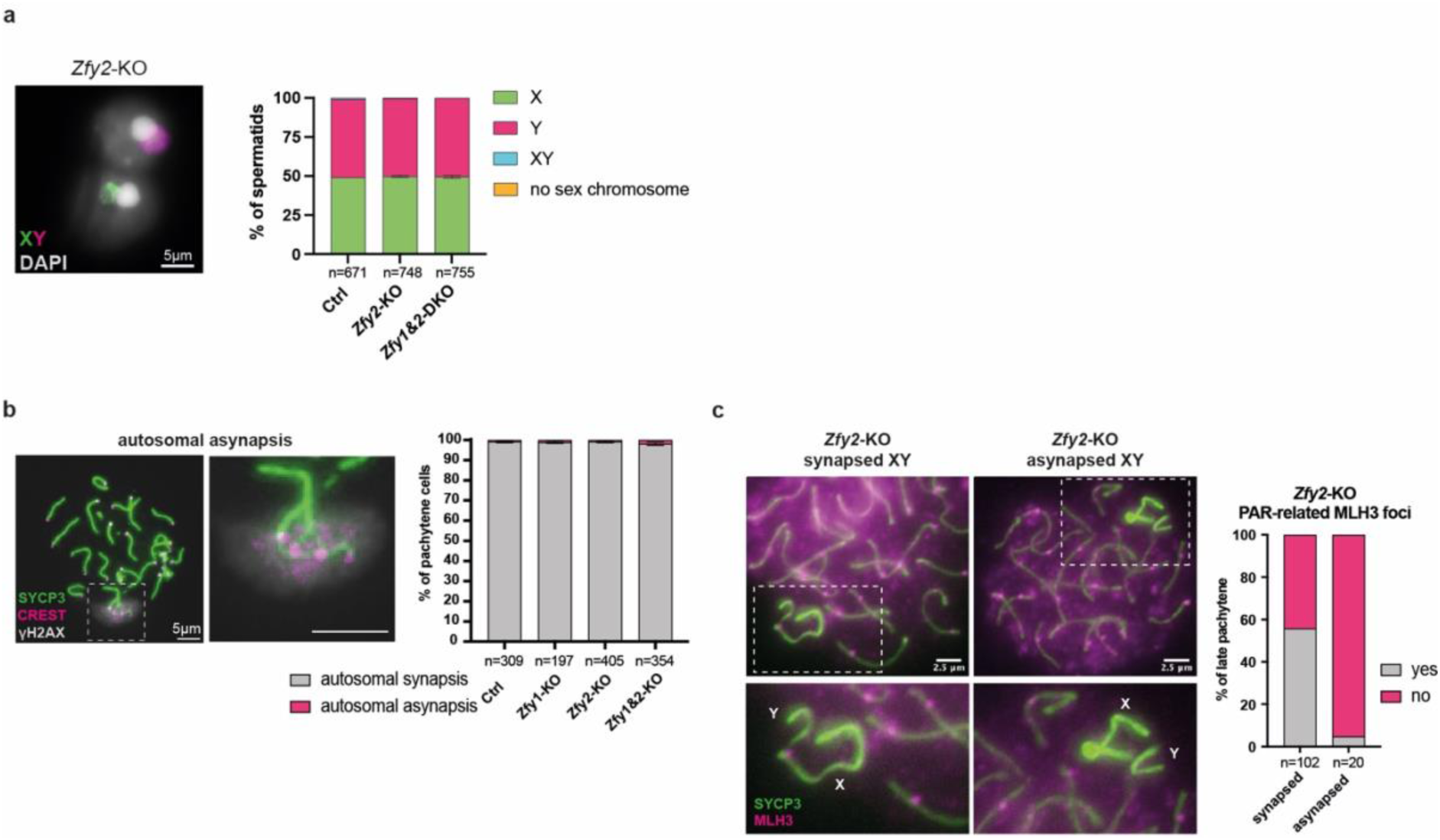
Characterisation of *Zfy* functions in chromosome synapsis. **a**, X and Y chromosome painting of round spermatids to quantify the percentage of spermatids bearing X, Y, XY or no sex chromosomes in controls, *Zfy2*-KO and *Zfy1&2*-DKO. Error bars = range. **b**, Pachytene spermatocytes immunostained for SYCP3 (green), CREST (magenta) and γH2AX (grey) to quantify autosomal asynapsis in control and *Zfy* deletants. Inset show an asynapsed autosome pair. n = number of pachytene counted across three biological replicates. Error bars = standard error of the mean. **c**, Late pachytene spermatocytes of *Zfy2*-KO immunostained for SYCP3 (green) and MLH3 (magenta) to quantify crossovers in sex chromosomes. n = number of cells counted across two biological replicates.

**Extended data Fig. 6:**
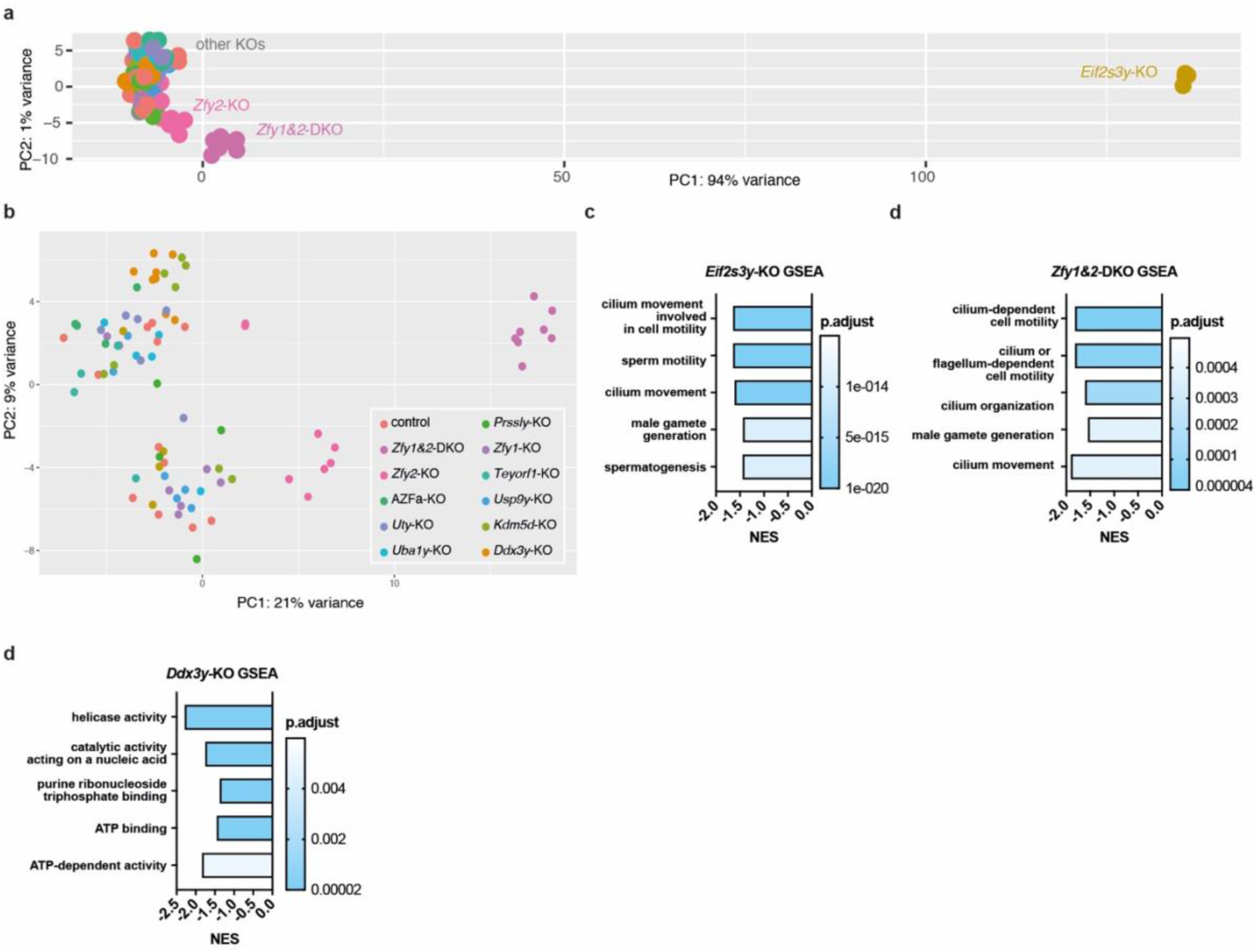
Characterising the effects of Y deletions on the testis bulk transcriptome. **a**, Principal component analysis (PCA) of testis bulk RNAseq for controls and the 13 Y-deletants. **b**, PCA excluding *Eif2s3y*-KO samples. **c-d**, Gene set enrichment analysis (GSEA) showing the top five downregulated gene ontology terms based on adjusted p-values in bulk samples of *Eif2s3y*-KO, *Zfy1&2*-DKO and *Ddx3y*-KO. The normalised enrichment score (NES) is plotted.

**Extended data Fig. 7:**
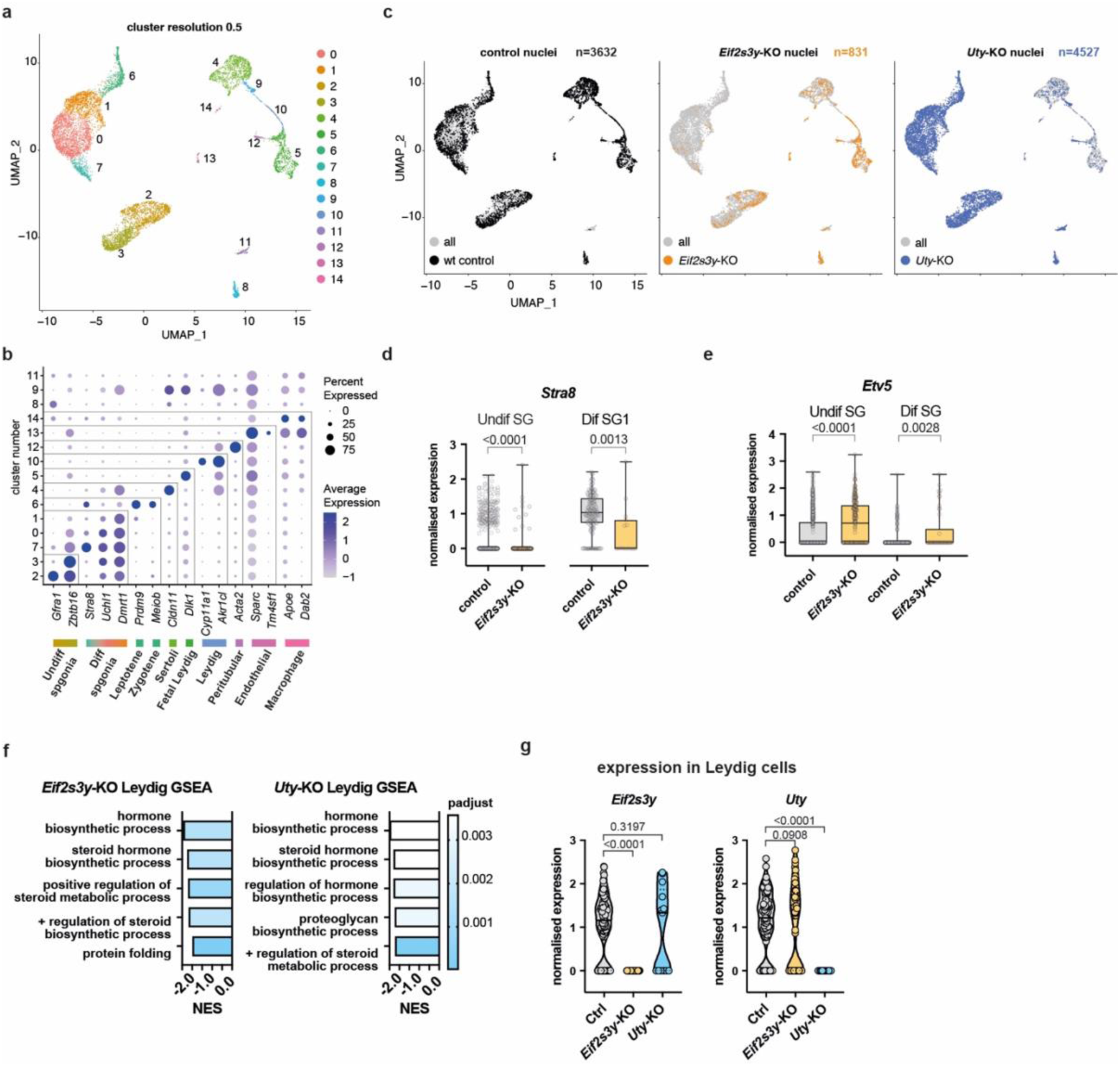
Single nuclei RNAseq of P10 control, *Eif2s3y*-KO and *Uty*-KO. **a**, Integrated UMAP of P10 testes clustered at the 0.5 resolution showing 15 clusters. **b**, Dot plot used for cell type assignment based on marker gene expression. Clusters 8, 9 and 11 do not have a clear signature and were annotated as “unknown”. **c**, UMAPs showing the contribution of each genotype. The number of nuclei is shown for each genotype. **d**, Normalised expression of the differentiation marker *Stra8* in merged undifferentiated (Undif SG) and differentiating spermatogonia 1 clusters (Dif SG1) for *Eif2s3y*-KO and control. **e**, Normalised expression of the progenitor marker *Etv5* in merged undifferentiated (Undif SG) and merged differentiating spermatogonia clusters (Dif SG) for *Eif2s3y*-KO and control. **f**, GSEA analysis in Leydig cells of *Eif2s3y*-KO and *Uty*-KO showing the top five downregulated gene ontology terms based on normalised enrichment score (NES). **g**, Expression of *Eif2s3y* and *Uty* in control, *Eif2s3y*-KO and *Uty*-KO. All p values calculated by Kolmogorov-Smirnov test.

**Extended data Fig. 8:**
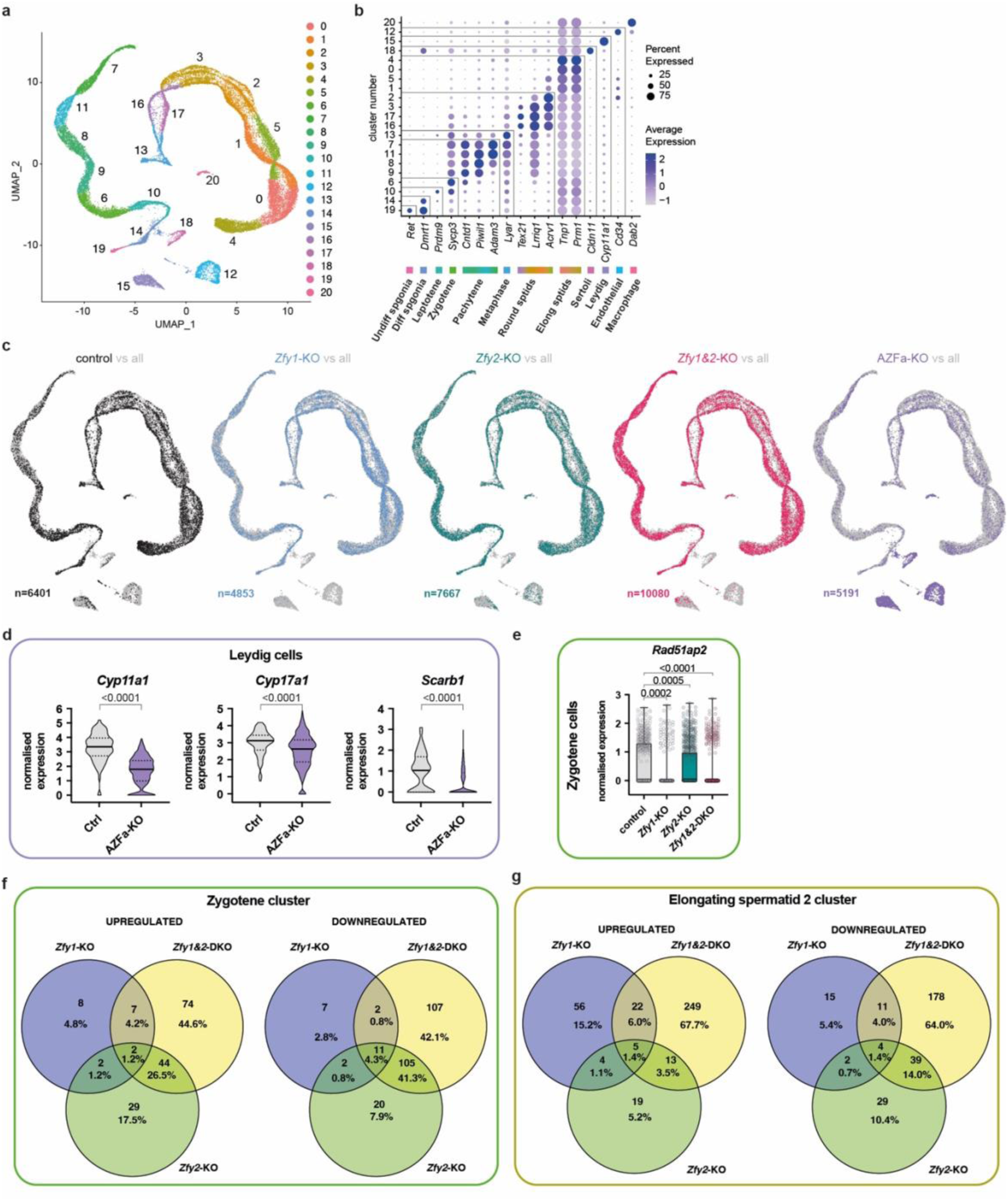
Single nuclei RNAseq of adult controls, *Zfy1, Zfy2, Zfy1&2* and AZFa KOs. **a**, Integrated UMAP of adult testes clustered at the 0.5 resolution showing 21 clusters. **b**, Dot plot used for cell type assignment based on marker gene expression. **c**, UMAPs showing the contribution of each genotype. The number of nuclei is shown for each genotype. **d**, Normalised expression of genes involved in the steroidogenesis pathway in adult control and AZFa-KO Leydig cells. **e**, Normalised expression of the recombinase *Rad51ap2* in control, *Zfy1*-KO, *Zfy2*-KO and *Zfy1&2*-DKO zygotene cells. **f-g**, Overlap in DE genes between *Zfy1*, *Zfy2* and *Zfy1&2* KOs in e, zygotene and f, elongating spermatid clusters.

## Methods

### Mouse colony maintenance

Mouse lines (*Mus musculus*) were maintained under UK Home Office Regulations, UK Animals (Scientific Procedures) Act 1986, and according to ethical guidelines at the Francis Crick Institute. Permission for animal experiments was granted by The Crick Biological Research Facility Strategic Oversight Committee (BRF-SOC) incorporating Animal Welfare and Ethical Review Body (AWERB) (Project Licence P8ECF28D9). Mice were kept as previously described^49^. Generated Y-deletant F0s were backcrossed to C57BL/6J females. For *Eif2s3y*-KO and *Zfy1&2*-DKO analyses, only F0 animals were used, as they could not reproduce. For the other genotypes including controls, F1 animals were used, except for IVF assays and litter size/sex ratio analyses, for which later (more backcrossed) generations were also used. Adult males were processed at ages between 13-15 weeks.

### mESC derivation and maintenance

We wanted the mESC line used for targeting to satisfy three requirements, 1) it should carry a reporter for easy quantification of ESC contribution in founder, 2) the Y chromosome should originate from a C57BL/6J background, and 3) it should possess high developmental potential. First, a B6D2F1-Tg(CAG/Su9-DsRed2, Acr3-EGFP RBGS002Osb)^50^ female was crossed to a C57BL/6J male and the line backcrossed to C57BL/6J for six generations. A resulting male carrying the transgenes was crossed to a CBA female, and the resulting blastocysts used for mESC derivation as previously described ^49^. The resulting mESCs were therefore CBAB6F1, with a C57BL/6J Y chromosome and carrying a mitochondrial DsRed2 and acrosomal GFP reporters. All mESC lines were maintained in 2i/LIF conditions on laminin-coated plates as previously described^51,52^.

### Preparation of the CRISPR-Cas9 components

sgRNAs were designed using the online tool CRISPRdirect^53^. sgRNAs with predicted low off-target activity were chosen (Supplementary Table 16). Additionally, for each selected sgRNA, potential off-target sites were identified using the Cas-OFFinder tool^54^ (Supplemetary Table 17). sgRNA oligonucleotides (synthesised by Eurofins Genomics, Germany) were annealed, and then ligated into the targeting plasmid px333-puro at the *BbsI* or *BsaI* (NEB, USA) restriction enzyme sites. The px333-puro vector was engineered by cloning a puromycin resistance cassette from px459v2 (gift from Feng Zhang, Addgene #62988) into the px333 vector (gift from Andrea Ventura, Addgene #64073) using standard directional cloning with *FseI* and *NotI* restriction enzymes (NEB, USA) (Extended data Fig.1b).

### Generating the Y-deletant mESCs

150,000 cells were seeded onto a laminin-coated well of a 6-well plate. On the following day, mESCs were transfected. First, 2.5μg of px-333-puro plasmid (encoding sgRNAs) was added to 100μl of Opti-MEM (ThermoFisher Scientific, USA), followed by 10μl of FuGENE® HD (Promega, USA) reagent. The solution was then mixed, incubated at room temperature for 10 minutes and then pipetted onto the cells. Two days post transfection, cell medium was changed to 2i+LIF containing puromycin (1.8μg/ml). Cells were kept in selection media for 48h, then put back into 2i+LIF for 4 days of recovery (Extended data Fig.1b). Individual surviving colonies were picked manually into the 96-well format and expanded. DNA extracted from individual clones was used for PCR screening. PCR reactions were carried out using the Q5 High-Fidelity DNA polymerase kit (New England Biolabs (NEB), USA) in a total volume of 25μl following manufacturer’s instructions. Primers (Supplementary Table 16) were designed using Primer3 (http://bioinfo.ut.ee/primer3/), synthesised by Eurofins Genomics (Germany), and used at the final concentration of 10μM. Agarose gel electrophoresis was used to confirm correct PCR amplification. Clones with a detectable deletion band and no exonic and boundary bands were amplified, frozen down and submitted for MiSeq and low pass WGS (Extended data Fig.1a, d, Supplementary Table 18).

### Tetraploid aggregations

2-cell embryos from plugged superovulated CD-1 mice were harvested and transferred into a culture dish with potassium supplemented simplex optimised medium (KSOM) micro drops. They were then transferred to Flushing and Holding Media (FHM), then to 0.3M mannitol. The embryos were then aligned into the fusion chamber containing mannitol, in between CUY5001P1 fusion electrode (Nepa Gene, Japan). The embryos were then pulsed using the Nepa ECFG21 (Nepa Gene, Japan) with a 10-volt 30 second AC pulse, followed by a 150-volt 30μs DC pulse, with one 0.1s interval and 10% decay and positive polarity. Embryos were then washed through three drops of FHM, a dish of KSOM, and cultured in a dish. Embryos that had successfully fused one hour post pulse were cultured until day 2. Acid Tyrode’s solution (Sigma, T1788) was used to remove the zona pellucida and the 4-cell stage embryos were then washed in 3 drops of FHM. Denuded embryos were washed in KSOM and individually placed in aggregation wells. Clumps containing 8-15 mESCs in KSOM were then added to the aggregation wells containing half of the individual embryos. The remaining half were individually deposited into the aggregation wells, “sandwiching” the mESCs in between embryos. Aggregated embryos were cultured overnight. On day 3, aggregated embryos at the blastocyst stage were washed and transferred into E2.5 pseudopregnant CD-1 females via uterine transfer. Overall, we achieved an average birth rate of 15.47% and average survival rate to adulthood of 85.39%.

### IVF assays

On day one, sperm was harvested by nicking the apex of one cauda epididymis and transferring the sperm into a 90μl drop of TYH+MBCD. The sperm solution was incubated for 30 minutes at 37°C. In the meantime, oocyte from B6CBAF1 females (superovulated using inhibin antiserum) were collected and incubated into 200μl drops of fertilisation medium (0.25mM glutathione in Human Tubal Fluid medium (HTF)) for maximum 30 minutes. 3-5μl of sperm was added into the fertilisation medium drops containing oocytes. 3 to 4 hours later, oocytes were washed in 4 drops of HTF, and the presumptive zygotes cultured overnight. On day two, the percentage of 2-cell embryos was assessed.

### Testis histology

Mouse testes were dissected, weighed, and fixed in Bouin’s (Sigma, UK) overnight at room temperature. The next day, they were washed in distilled water and stored in 70% EtOH at room temperature. Fixed testes were embedded in wax, sectioned transversely at 4μm and stained with Periodic Acid-Schiff (PAS). Sections were imaged using an Olympus VS120 Slide Scanner with a 40X objective.

### Immunostaining of testis sections

Testes were fixed in 4% paraformaldehyde (PFA) overnight at 4°C, washed in 70% EtOH, embedded in paraffin and sectioned at 4μM on slides. Slides were incubated at 60°C for 10 minutes, then washed in Histo-Clear (National Diagnostics, USA) twice for 5 minutes. Samples were then rehydrated through ethanol washes (2x 100% ethanol 1 minute, 2x 95% ethanol 1 minute, once in 75% ethanol 1 minute and once in distilled water 1 minute). Antigen retrieval was performed by boiling the slides in 0.01M Sodium Citrate for 10 minutes and then letting the solution cool for 15 minutes at room temperature. Sections were blocked in 5% FBS in PBS with 0.01% Triton at room temperature for one hour. For apoptosis quantification, slides were incubated with rabbit anti-cleaved poly-ADP-ribose polymerases (cPARP), (1:200, ab32064, Abcam) in a humidified chamber at 37°C for 2 hours. Samples were incubated in the secondary AlexaFluor 647 anti-rabbit antibody (1:200, ThermoFisher) at room temperature for 1 hour. For spermatogonia quantification, slides were incubated with LIN28A (1:200, ab63740, Abcam) and goat cKIT (1:100, AF1356, R&D systems) at 4°C overnight. Samples were incubated in the secondary AlexaFluor 647 anti-goat and AlexaFluor 594 anti-rabbit antibodies (1:250, ThermoFisher) at room temperature for 1 hour. Washes were performed with 0.01% PBS-Triton. Sections were counterstained with the DNA-binding fluorescent stain DAPI (D9564, Sigma-Aldrich).

### Immunostaining of testis nuclear spreads

Nuclear spreads were prepared from frozen samples, except for MLH3 quantification, where fresh tissue was used. For frozen samples, a piece of testis was thawed in a drop of PBS and chopped into a suspension using 2 scalpels. One drop of the testis suspension was dropped onto a pre-boiled slide from 15-20 cm height. One drop of 0.05% Triton X-100 (dissolved in distilled water) was added to each drop and left for 10 minutes. 8 drops of 2% formaldehyde, 0.02% SDS in PBS were then added and slides left incubating at room temperature in a humidified chamber for one hour. Slides were then quickly dipped in distilled water 6 times and aired dried for 5 minutes. Slides were either used for immunostaining straight away or stored at −80°C.

Samples from fresh tissues were prepared by adapting a previously published protocol^55^. A small piece of testis was incubated in 2.5mL dissociation buffer (0.25% Trypsin, 20μg/μl DNAse I in PBS) at 37°C for 15min shaking at 250rpm. The tissue was further dissociated by pipetting for ∼2min using a wide-mouthed Pasteur pipette and the reaction quenched by adding 150μl of FBS. Cells were moved through a 70μm cell strainer and pelleted by spinning at 1000g for 5min. The pellet was resuspended in 15mL of PBS with 20μg/ml DNAse I the further spun down at 1000xg for 5min. This was repeated for a total of 3 washes and cells finally resuspended in PBS (∼200μl of PBS per 100mg of tissue). 10μl of suspension was added into a tube containing 90μl of 75mM sucrose in water. 45μl of the dissociated cells was added to a slide, inside a square drawn with a lipophilic marker containing 100μl fixing solution (1% PFA, 0.15% triton, in PBS). Slides were incubated in a humidified chamber overnight. Slides were dried, rinsed in dH2O, washed 1min in PBS 0.4% Kodak PhotoFlo 200, air-dried and stored at −80°C.

For immunostaining, slides were incubated in PBT (PBS, 0.15% BSA, 0.1%Tween-20) for one hour at room temperature. The primary antibodies diluted in PBT were then added to the slides and incubated overnight at 37°C in a humidified chamber. The following antibodies were used: guinea pig SYCP3 (in house) at 1:100, human CREST (gift from W. Earnshaw) at 1:100, mouse γH2AX (05-636; RRID: AB_309864, Millipore) at 1:250, rabbit MLH3 (gift from Paola Cohen) at 1:500. The slides were washed in PBS for 5 minutes 3 times. The secondary antibodies (Alexa Fluor® 488, Alexa Fluor 594 and Alexa Fluor 647) diluted in PBS (1:250), were then added to the slides, and incubated for 1 hour at 37°C. Slides were washed for 5 minutes 3 times and mounted in Vectashield plus DAPI (4’,6-diamidino-2-phenylindole) (Vector Laboratories, USA). Stained testis spreads were imaged using an Olympus delta vision or Zeiss Observer microscope using 40X or 63X objectives.

### Chromosome painting

Testis nuclear spreads or testis sections (see previous two sections) were washed twice in PBS for 5 minutes and dehydrated through ethanol series, 70% ethanol twice, 90% ethanol twice, 2 minutes each and 100% ethanol once for 5 minutes. Slides were then washed three times in 2xSSC for 5 minutes and denatured in 2xSSC at 80°C for 5 minutes. Slides were quenched in ice-cold 70% ethanol and a second dehydration step performed as described above. The X (XMP X green, D-1420-050-FI) and Y (XMP Y orange, D-141421-050-OR) chromosome paints (Metasystem, Germany) were added to the slides in equal volumes (10-20μl total), a coverslip added, and the slides incubated, first at 75°C for 5 minutes and then at 37°C overnight in a humidified chamber. Slides were washed in 2xSSC at room temperature, then in 0.4xSSC at 72°C for 2 minutes, then in 2xSSC with 0.05% Tween-20 at room temperature for 30 seconds. Slides were rinsed in distilled water and stained with DAPI for 10 minutes.

### Sperm morphology analyses

To extract spermatozoa, three incisions were made on the cauda epididymis, which was then dropped into a tube containing 200μl of pre-warmed TYH+MBCD and incubated in a 37°C 5% CO2 incubator for 8 minutes. Using tweezers, the cauda was then removed from the tube and additional 200-400μl of warm TYH+MBCD was added. After gently mixing by pipetting with a wide-bore tip, 50μl of sperm solution was transferred drop-by-drop to a new tube containing an equal volume of 4% PFA (i.e., final concentration of 2% PFA). After 5 minutes of incubation at room temperature, spermatozoa were spun down at 4°C for 3 minutes at 300g. The sperm pellet was resuspended in a staining solution (0.1% polyvinyl alcohol (PVA), Hoechst (1:1000, ThermoFisher, 62249), Phalloidin (1:250, ThermoFisher, 12381), Triton (0.1%, Sigma, 93443) in PBS). The sperm solution was then pipetted onto a SUPERFROST slides (Fisher Scientific, UK) (15μl per slide) and kept sealed at 4°C. With a Nikon LTTL 3 microscope, micromanager and a Python script, a strategy to automatically map and image individual sperm heads was designed. First, slides containing fixed spermatozoa were scanned at the 20X magnification on the DAPI channel and positions of individual sperm heads were recorded. Switching to a Nikon Plan Apo 100x/1.4 objective, the saved locations (sperm heads) were then automatically imaged with a z-stack. Z-stacks were then analysed using an ImageJ script to identify the best focused image in the stack and to normalise the exposure settings across files. At least 100 sperm heads per animal were imaged in that way. Sperm head images were imported into the Nuclear_Morphology software for morphology analysis^56^. Analysis settings were left to default except for “flattening threshold” which was changed to “120”. After detection, identified sperm heads were manually curated to exclude any poorly outlined sperm heads from the analysis.

### Sperm motility analyses

After isolation (see sperm morphology analysis), 3μl of sperm solution was loaded into a Leja 20μm 4 chamber slide (IMV Technologies, France). The slide was kept on a portable warming stage (HS-PT-USB, Microptic, Spain) set to 37°C. Once on the slide, the sperm solution was let to settle for 2 minutes before analysis. Altogether, sperm motility analysis was performed using the SCA Motility and concentration TOX edition (SCA-TOX-01, Microptic, Spain). Analysis started 15 minutes after dissection of the cauda epididymis. Set parameters were used and at least 20 video fields captured throughout the slide to obtain data for at least 300 spermatozoa. After capture, each field was inspected manually to delete any sperm-tracking errors or duplicated sperm tracks.

### Sperm counts

Cauda epididymis were dissected and shredded in pre-warmed TYH+MBCD medium (usually 1.1ml) to extract all the sperm. Samples were incubated in a 37°C 5% CO2 incubator for up to 30 minutes. 3μl of sperm solution was loaded into a Leja 20μm 4 chamber slide and the total cauda sperm count was estimated using the SCA software.

### MiSeq

MiSeq was used to characterise on-target deletion outcome and to screen for potential off-target editing. PCR amplicons were purified using SPRI bead clean, and library prepared according to the Illumina MiSeq library prep manufacturer’s instructions (Nextera Index Kit V2). Samples first underwent PCR indexing and libraries were then normalised and pooled after fluorometric quantification using a dsDNA dye. Library were sequenced using the Illumina MiSeq platform with a PE 250bp run configuration on a Nano flowcell. An average of 2000-5000 reads were obtained per samples. After sequencing, the Fastq sequence files were collapsed using the FastX Toolkit (v0.0.13) [https://github.com/agordon/fastx_toolkit]. To determine the whole-locus deletion outcome, the collapsed MiSeq reads were aligned to the reference mouse genome (*Mm10*) using *blastn* (Extended data Fig.1c, g). To examine potential off-target editing, the collapsed MiSeq reads were aligned using Burrows-Wheeler Alignment tool (BWA, v0.7.170)^57^ and analysed using the R package CrispRVariants^58^ (Extended data Fig.1h). This was used to calculate the proportion of wild type reads (perfect match to *Mm10*) around the off-target site compared to the sum of all other reads containing single nucleotide variants ± 20bp away from the sgRNA binding site.

### Low pass whole genome sequencing

Low pass WGS was used to confirm whole-locus deletion and to ensure that the rest of the Y chromosome, as well as the X homologues were intact. It also allowed to characterise the karyotype of each cell line. After genomic DNA purification, samples were prepared with the Illumina DNA Prep with Enrichment kit following the Nextera flex protocol. Coverage of 0.1X-0.3X was obtained on average. Resulting FastQ reads were aligned to the reference genome using BWA (Extended data Fig.1e). The R package QDNAseq^59^ was used for karyotype analysis (Extended data Fig.1f).

### Bulk RNAseq

About 20mg of frozen testis pieces were placed in Precellys Homogenizer tubes 2.8 mm ceramic reinforced (Precellys 50722019, Bertin Technologies, France) on dry ice. 500μl of Trizol reagent (Sigma-Aldrich, USA) was added to the tubes and the samples placed in a Bertin Precellys 24-Dual High-throughput Tissue Homogenizer (Bertin Technologies, France). The machine was then run with the factory programme 5: 6,500 rpm for 20 seconds, automatic 25 seconds of pause and a repeat of 6500 rpm for 20 seconds. The lysate was then centrifuged for 5 minutes at 12,000 x g at 4°C and the clear supernatant transferred to a MaXtract High Density phase lock tube (Qiagen, UK). After 5 minutes of incubation at room temperature, 100μl of chloroform was added and tubes shaken vigorously for 2-3 minutes. Tubes were spun down for 15 minutes at 12,000 x g at 4°C. The upper aqueous phase was transferred to a new tube and 250μl of isopropanol added and mixed by pipetting. After 10 minutes of incubation at room temperature samples were spun down for 10 minutes at 12,000 x g at 4°C. The supernatant was discarded, the RNA pellet washed with 1ml of 75% ethanol and the samples spun down for 5 minutes at 7,500 x g at 4°C. Finally, the pellet was air-dried for 5-10 minutes and resuspended in 40μl of nuclease-free water. Samples were checked using a TapeStation (Agilent Technologies, USA). Libraries were prepared using KAPA mRNA HyperPrep Kits according to manufacturer instructions (KAPA Biosystems, Roche). Raw RNA-seq reads were processed using the RNA-seq nf-core pipeline (v3.2), star_rsem was used to generate raw reads counts. The read counts were processed in R using the DESeq2 (v1.34) package. Very lowly expressed genes were filtered out by applying a rowSums filter of >=5 to the raw counts table. Principal component analysis (PCA) plots were generated using the top 500 most variable genes, applying the vst() function and limma::removeBatchEffect() to remove batch effects for PCA visualisation. Raw counts were batch corrected and normalised using the DESeq() function, specifying “∼batch + Sample_Genotype” in the design formula. Log2 fold change (log2FC) and adjusted p-values between KO and WT were calculated using the results() function in DESeq2, specifying the “BH” (Benjamini-Hochberg p-value adjustment). Log2FCs were shrunk using the lfcShrink(type=”apeglm”) function. Differentially expressed genes (DEGs) were defined as genes which had an adjusted p-value < 0.05 and log2FC > 0.5 (Supplementary Table 1). Enrichment of genomic functions and cellular processes was done using the gseGO() function, as part of the R package, clusterProfiler (v4.2.2) (Supplementary Table 2).

### Single nuclei RNAseq

For *Eif2s3y* and *Uty* deletants, P10 testes were used because, unlike adult testes, they are enriched in spermatogonia, the cell type affected in these mutants. Moreover, for *Eif2s3y*, we did not want to capture potential secondary effects of the total spermatogonia arrest. For *Zfy2*, *Zfy1*-KO, *Zfy1&2*-DKO and AZFa-KO, which have all germ cell types, we sequenced nuclei from adult testes.

Testis nuclei isolation was adapted from a previously published protocol^36^. On ice, around 10mg of frozen testis was homogenised in 1ml of lysis buffer (250 mM sucrose, 25 mM KCl, 5 mM MgCl2, 10 mM HEPES pH 8, 1% BSA, 0.1% IGEPAL and freshly added 1 µM DTT, 0.4 U µl−1 RNase Inhibitor (New England BioLabs), 0.2 U µl−1 SUPERasIn (ThermoFischer Scientific)) and incubated on ice for 4 minutes. The lysate was filtered through a 30µm filter and then centrifuged at 500g for 5 minutes at 4C. The supernatant was removed, and the pellet resuspended in 1ml of wash buffer 1 (250 mM sucrose, 25 mM KCl, 5 mM MgCl2, 10 mM HEPES pH 8, 1% BSA, and freshly added 1 µM DTT, 0.4 U µl−1 RNase Inhibitor, 0.2 U µl−1 SUPERasIn). This was spun down at 500g for 5 minutes at 4°C and washing step repeated. Next the pellet was resuspended in 670 µl of freshly diluted dithio-bis (succinimidyl propionate) (DSP) (1mg/ml in PBS) and incubated at room temperature for 30 minutes. Fixation was quenched by adding 14.07µl Tris-HCL, pH8. Nuclei were spined down at 500g for 5 minutes at 4C and resuspended in 1ml of wash buffer 2 (250 mM sucrose, 25 mM KCl, 5 mM MgCl2, 10mM Tris-HCL, pH8, 1% BSA, and freshly added 1 µM DTT, 0.4 U µl−1 RNase Inhibitor, 0.2 U µl−1 SUPERasIn). The sample were spun down one more time at 500g for 4 minutes at 4°C and the pellet resuspended in 0.04% BSA in PBS.

Replicates from the P10 and adult 10x nuclei datasets were aligned to the mm39 genome using STARsolo (v2.7.9), specifying the parameters –soloMultiMappers EM –outFilterScoreMin 30 --soloFeatures Gene GeneFull --soloType CB_UMI_Simple --soloFeatures Gene GeneFull –soloUMIlen 12 --soloCBmatchWLtype 1MM_multi_Nbase_pseudocounts --soloUMIdedup 1MM_CR. Empty droplets were separated from nuclei using the proportion of reads mapping to intronic reads (Supplementary Table 19). Replicates were then processed in the Seurat R package (v4.3.0). Cells were retained if they had <5% reads aligning to mitochondrial and rRNA genes, if the total number of molecules within a cell was between 1000 and 50000 and the number of genes detected per cell was between 500 and 10000. The data was normalised using NormalizeData, then FindVariableFeatures. The dataset was scaled using ScaleData. Principle Component Analysis was applied using RunPCA followed by FindNeighbours and FindClusters. Uniform Manifold Approximation and Projection was estimated for the integrated dataset using RunUMAP. Doublets were identified using the DoubletFinder R package (v2.0.3) (Supplementary Table 19). Replicates from each KO were integrated together, then the integrated KOs from their respective P10 and adult datasets were integrated with each other using SelectIntegrationFeatures, FindIntegrationAnchors and IntegrateData, generating one integrated dataset for the P10 10x data, and another for the adult 10x data. Each dataset was processed using ScaleData, then RunPCA and RunUMAP. Clusters were identified using a 0.5 resolution for both the adult and P10 datasets. In the adult dataset, ambient RNA clusters were identified using the FindAllMarkers function, and subsequently removed. Clusters were annotated using published marker genes^36,37^. Differentially expressed genes were identified between wildtype control and KOs on a cluster-by-cluster basis using FindMarkers (Supplementary Tables 3 to 8). Enrichment of genomic functions and cellular processes was done using the gseGO() function, as part of the R package, clusterProfiler (v4.2.2) (Supplementary Tables 9 to 14).

## Supplementary discussion

Our chromosome painting analysis did not find increased rates of XY aneuploidies in *Zfy2*-KO, suggesting that the SAC sensitivity is unaltered. This contrasts with previous research that showed that addition of a *Zfy2* transgene in XO models re-established the elimination of spermatocytes with unpaired chromosomes^18^. However, it is essential to recognise the disparities between the experimental conditions employed in the cited study compared to ours. Vernet and colleagues employed transgene rescue techniques in which the expression levels of *Zfy2* exhibited notable deviations from the wildtype context^17,18^. This could have direct or indirect effects on SAC activation. Additionally, Vernet and colleagues used XO models, in which only one chromosome is misaligned at metaphase. In contrast, in this study, most of the Y, including the PAR remains intact, meaning that when X and Y fail to pair, two chromosomes are misaligned. This could trigger a more robust signal for SAC-mediated apoptotic elimination compared to XOs. If *Zfy2* acts to “enhance” SAC response, it may be needed to eliminate cells that have only one misaligned chromosome, and therefore low levels of SAC activation. In cells with more than one misaligned chromosome and therefore stronger SAC activation, *Zfy2* may be dispensable for apoptosis.

## Author contributions

J.M.A.T. and J.S. conceived the project. J.S., I.B.B. and M.B. generated the Y-deletant mESCs. J.S. performed testis weight, sperm count, sperm head and sperm motility analyses. J.S. extracted bulk RNA and single nuclei for RNA sequencing. J.S. and D.d.R. analysed testis histology. J.S., I.B.B., and G.S. performed and analysed immunostaining. I.B. performed XY chromosome painting. O.O. maintained the mouse colonies. W.V. performed analysis of bulk and snRNAseq datasets. V.M. derived the initial mESC line and advised on immunostaining and nuclei extraction. B.G. advised on snRNAseq analysis. K.C. and K.C. oversaw the tetraploid aggregations. J.S. and J.M.A.T. wrote the manuscript.

## Acknowledgments

The authors thank the Francis Crick Institute Genetic Modification Service (GeMS), Biological Research, Experimental Histopathology, Advanced Light Microscopy and Advanced Sequencing facilities for their contributions and expertise. We thank the members of the J.M.A.T. lab for comments and discussion on the manuscript. This work was supported by the Francis Crick Institute which receives its core funding from Cancer Research UK (CC2052), the UK Medical Research Council (CC2052), and the Wellcome Trust (CC2052).

